# Predicting Discrete Structural Transformations in Small Molecules from Tandem Mass Spectrometry

**DOI:** 10.64898/2026.05.06.723373

**Authors:** Xianghu Wang, Gwendolyn Kiler, Daniela Herrera-Rosero, Mohammed Reza Shahneh, Michael Strobel, Christian Geibel, Yasin El Abiead, Vanessa V. Phelan, Daniel Petras, Mingxun Wang

**Affiliations:** Department of Computer Science and Engineering, University of California, Riverside, Riverside, CA, United States; Department of Pharmaceutical Biology, University of Tübingen, Tübingen, Germany; Department of Microbial Bioactive Compounds, University of Tübingen, Tübingen, Germany; Department of Natural Sciences and Sustainable Resources, Institute of Analytical Chemistry, BOKU University, Vienna, Austria; Department of Pharmaceutical Sciences, University of Colorado Anschutz Medical Campus, Aurora, CO, United States; Department of Biochemistry, University of California, Riverside, Riverside, CA, United States

**Keywords:** Tandem mass spectrometry, Metabolomics, Machine learning, Structure elucidation, Molecular annotation, Structural transformation prediction

## Abstract

Tandem mass spectrometry (MS/MS) fragments molecules into smaller pieces, generating spectra composed of m/z values and intensities that encode structural information for molecular annotation. With increasing mass spectrometry data acquisition speeds, manual annotation from MS/MS lags far behind data generation and remains a bottleneck in metabolite annotation. Current computational methods, such as molecular networking, address this challenge by organizing similar structures into families of related compounds. However, they generally provide only similarity scores, offering weak actionable insights for structural annotation. To address this limitation, we present the Molecular Transformation Graph Edit Measure (MT-GEM), a distance metric that quantifies discrete structural transformations between molecules through graph edge removals that approximate structural modifications. Building on this metric, we developed an ensemble machine learning architecture, the Spectrum Transformation Edit Predictor (STEP), that builds upon TransExION and DREAMS to predict MT-GEM distances from MS/MS spectra. STEP achieves an average precision of 48.4% for identifying single structural transformations between MS/MS pairs, representing more than a tenfold improvement over state-of-the-art similarity metrics, including spectral entropy similarity (3.8%) and modified cosine (2.5%). On experimental human gut microbial community data, STEP identifies 3 times more single-transformation metabolite pairs than feature-based molecular networking at equivalent precision. In a discovery application, STEP highlights one drug metabolite and two new natural product analogs missed by modified cosine in feature-based molecular networking. By providing discrete transformation predictions rather than continuous similarity scores, MT-GEM and STEP enable hypothesis-driven metabolite annotation with testable structural modifications, which we envision will accelerate discovery of new molecules from MS/MS metabolomics datasets.

## Introduction

Untargeted tandem mass spectrometry (MS/MS) measures chemical diversity at scale, generating hundreds of spectra per second (1). However, data volume far exceeds manual annotation capacity. This bottleneck is consequential because MS/MS is central to modern metabolomics and natural product research. As a high-throughput analytical technique, MS/MS enables rapid screening of complex biological and environmental samples, identification of known compounds through spectral library matching, and detection of novel structural analogs related to known scaffolds, characterization of drug metabolites and biotransformation products, and assessment of chemical structural diversity across sample cohorts (2, 3). Each of these applications relies on relating experimental spectra to molecular structures, making the gap between data acquisition rate and annotation capacity a direct constraint on discovery throughput.

Three computational approaches attempt to address this bottleneck. MS/MS matching to spectral libraries compares unknown experimental spectra against reference MS/MS libraries (4). Given the success of these approaches, still 87% of MS/MS spectra remain unannotated (5, 6). Spectral similarity tools extend beyond exact matches in MS/MS library search by exploiting the fact that molecules with shared substructures tend to fragment similarly, producing related peak patterns in MS/MS. Modified cosine aligns shifted peaks between spectra to account for precursor mass differences (7), while spectral entropy compares the information content of combined versus individual spectra to quantify shared fragmentation (8). Machine learning (ML) models extend these MS/MS similarity approaches by predicting structural similarity directly from MS/MS data, often in the form of Tanimoto coefficients (9). MS2DeepScore (10) and TransExION (11) are two ML approaches that try to tackle this challenge. A key application of all these strategies is to propagate structural annotations from library-matched compounds to unannotated features, using spectral relationships to infer that an unknown compound is structurally related to a known reference and thereby narrow the space of candidate structures.

However, both spectral similarity and ML methods produce a continuous similarity score between two MS/MS spectra that yield weak actionable computational paths for solving the structure of unknown compounds. A predicted Tanimoto coefficient of 0.85 indicates similarity without revealing what differs, how many modifications separate the molecules, or where modifications occur, leaving chemists without testable structural hypotheses. Recent work has begun to address this limitation by defining discrete molecular distances. The Maximum Common Edge Substructure (MCES) distance counts the minimum number of individual bond edits (additions, deletions, or substitutions) required to transform one molecular graph into another (12). While MCES provides a well-defined structural distance, its bond-level granularity does not align well with how chemists can reason about structural modifications. For transformations involving the addition or removal of multi-atom functional groups at single bond cleavage sites, MCES counts edits for every bond within the added or removed group, while MT-GEM counts the entire group as a single substructure operation, aligning more closely with how chemists describe discrete structural modifications.

To better capture the granularity of chemical transformations, we introduce the Molecular Transformation Graph Edit Measure (MT-GEM). MT-GEM operates at the substructure level rather than the bond level. It identifies the maximum common substructure (MCS) between two molecules then counts the minimum number of connected components that must be removed from each molecule to isolate the MCS. Because MS/MS fragments molecules at chemical bonds, substructure removal in graphs approximates bond fragmentation in the mass spectrometer, providing an interpretable measure of chemical distance (13–16). An MT-GEM distance of 1, therefore, indicates that two molecules differ by exactly one such substructure addition or removal, representing a single discrete chemical transformation such as hydroxylation, methylation, or deglycosylation.

However, both MCES and MT-GEM calculations are NP-Hard, with computational time increasing exponentially with molecular size (17). More critically, both metrics require known molecular graphs and therefore cannot be applied to unannotated spectra, precisely the features that most need structural characterization. A predictive model (18) that estimates MT-GEM distances directly from MS/MS spectra would circumvent both limitations, enabling discrete transformation prediction without prior structural knowledge and at a fraction of the computational cost.

Here we introduce STEP (Spectrum Transformation Edit Predictor), an ensemble machine learning model that integrates TransExION and DreaMS (18), which is trained to predict discrete structural transformations directly from MS/MS spectra pairs, specifically MT-GEM distances of 1. By explicitly linking MS/MS spectra in this way, STEP enhances the ability to discover new molecules by connecting annotated library-matched MS/MS to putative structural analogs by prioritizing pairs most likely differing by a single structural transformation. By integrating downstream computational approaches, we can constrain the candidate modifications to specific functional groups (e.g., +15.995 Da suggests hydroxylation) and propose the site of modification with ModiFinder (19) on the known structural scaffold. Together, this creates a complete annotation propagation pipeline: STEP identifies the candidate pair, and ModiFinder pinpoints its location, converting an unannotated feature into a specific, testable structural hypothesis that accelerates manual MS/MS interpretation.

We found that STEP outperformed existing spectral similarity and machine learning baselines (20, 21). Applied to real-world experimental data from a synthetic gut-community drug metabolism study (22) and a bioactivity-guided natural product dataset (23), STEP identified substantially more single-transformation pairs than feature-based molecular networking (24) that as a demonstration, resulted in the discovery of three structural analogs.

## Results

### Data Preprocessing

We collected 1,320,389 MS/MS from NIST20 and 825,973 MS/MS from GNPS spectral libraries for a total of 2,146,362 spectra in the combined GNPSNIST dataset. We preprocessed and filtered GNPSNIST by keeping only positive mode MS/MS spectra in the precursor m/z range 10-1000 m/z with a minimum of 5 peaks, resulting in 490,292 MS/MS spectra and 32,695 structures. We truncated spectra to the top 100 most abundant peaks, applied a 3.0 m/z sliding window to remove dense peak clusters, normalized peaks from 0.0 to 1.0 relative to maximum intensity, and removed peaks below 0.1% intensity. This filtering resulted in 416,099 spectra representing 31,344 unique structures (see Methods **Data Acquisition and Preprocessing**).

To prevent data leakage, we employed the dataset splitting methodology from Strobel et al., partitioning by maximizing pairwise Tanimoto distance between molecular structures across train and test sets (25). A total of 500 structures were randomly selected as a holdout validation set from the training set. This partitioned the GNPSNIST dataset into 271,413 MS/MS from 21,365 unique structures for training (GNPSNIST-train), 51,183 MS/MS from 3,740 unique structures for testing (GNPSNIST-test), and 6,306 MS/MS from 500 unique structures for validation (GNPSNIST-validation). We calculated MT-GEM distances for all eligible MS/MS pairs from the GNPSNIST-train, validation, and test datasets. Because MT-GEM distance of one occurs rarely by random chance, we enriched for “distance one” pairs of MS/MS spectra to 2% (The natural prevalence of MT-GEM distance-one pairs in the GNPSNIST-train dataset was approximately 0.19%) of the pairs used in the GNPSNIST-train dataset. This produced 230 million pairs for training in GNPSNIST-train. For the GNPSNIST-test, we calculated 3,880,795 pairs of MS/MS spectra and their accompanying MT-GEM. The prevalence of MT-GEM “distance one” pairs was 0.3% (11,666 pairs), with 3,869,129 pairs with non-1 MT-GEM distance.

### Overall model performance on GNPSNIST

To measure the performance of the STEP model, we calculated the precision (fraction of predicted single-transformation pairs that are correct) and recall (fraction of actual single-transformation pairs successfully identified) of STEP against alternative approaches and comparable architectures by using the MS/MS similarity predictions as proxies for MT-GEM 1 confidence thresholds.

Across 3,880,795 MS/MS pairs (with 0.3% positive class prevalence) in GNPSNIST-test, we measured the performance under two evaluation schemes: spectrum-level, where each MS/MS pair prediction performance is measured, and structure-level, where precision and recall are averaged per unique structure pair. We measured under the additional structural evaluation scheme in addition to the common spectrum level evaluation as we noticed many high performance cases of duplicate structure pairs skewing model performance upwards, which gave a biased view of the model’s expected performance in discovering structural pairs (see Methods **Evaluation Protocol for GNPSNIST**). **Fig. 1** shows precision-recall curves for spectrum-level (**Fig. 1A**) and structure-level (**Fig. 1B**) performance, with STEP achieving 48.4% area under the precision-recall curve (AUPRC) versus MS2DeepScore’s 8.3%.

**Fig. 1.**
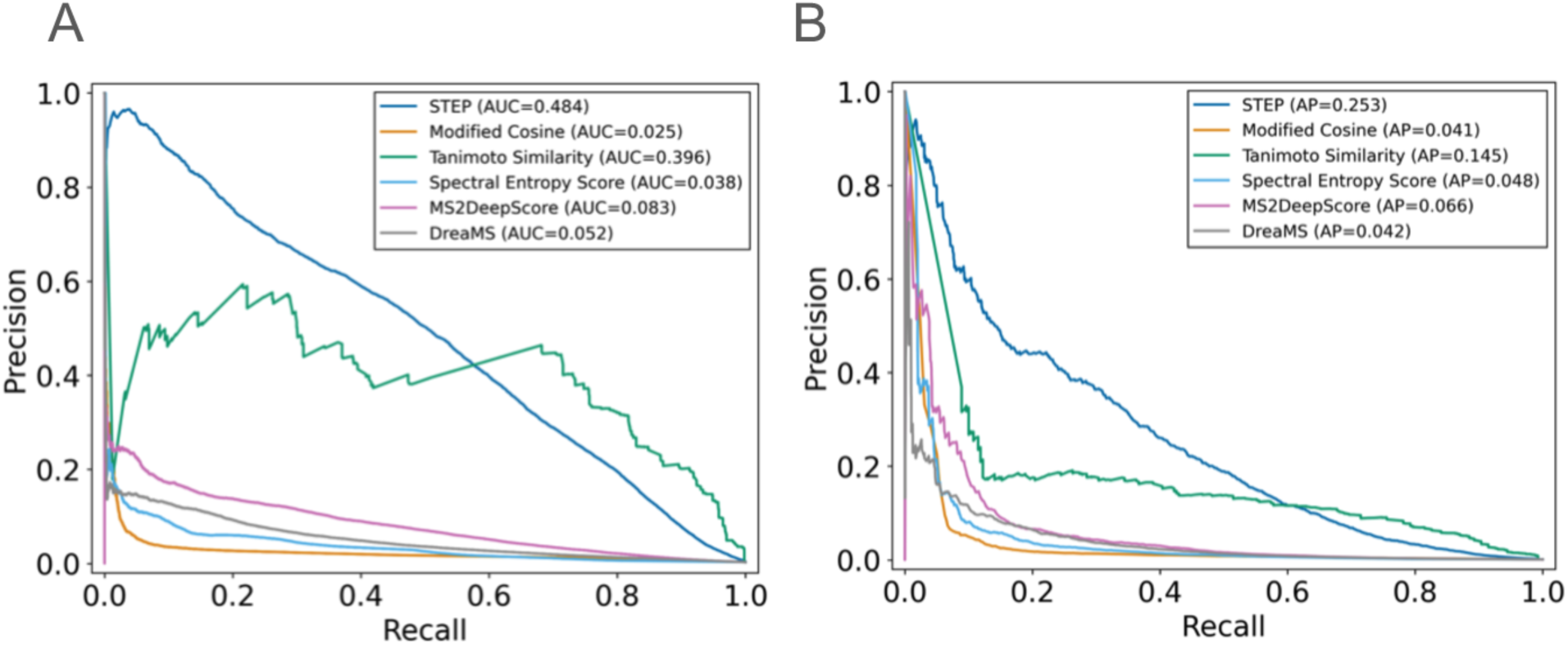
Benchmark performance of STEP on the GNPSNIST test set. (A) Precision recall curves depicting spectrum level model performance on the GNPSNIST test set. STEP (Blue with AUPRC 0.484) improves upon prediction performance compared to the next best method, MS2DeepScore (Pink, with AUPRC 0.083), as well as the metric that MS2DeepScore attempts to predict, Tanimoto Similarity (Green with AUPRC 0.396), with a more pronounced difference at higher levels of precision below 0.4 recall. (B) Precision recall curves depicting structure-level model performance on the GNPSNIST test set. When multiple spectrum pairs exist for each unique structure pair, the score contributions to precision/recall are averaged per unique structure pair to account for certain structure pairs having many MS/MS spectra. In the unique-structure metric, STEP exceeds the Area Under Precision Recall Curve (AUPRC) and precision of predicting MT-GEM Distance one at all confidence thresholds with recall <60%.

Under spectrum-level evaluation, at 5% recall, STEP achieved 94% precision compared to 22.4% for MS2DeepScore, representing a more than 4-fold improvement. At a fixed precision of 20%, STEP achieved 80% recall, whereas MS2DeepScore achieved 8% recall, representing a 10-fold improvement. Overall, STEP achieved an area under the precision-recall curve (AUPRC) of 48.4%, compared to 8.3% for MS2DeepScore, representing an over 5-fold improvement.

Under structure-level evaluation, reflecting on molecular diversity rather than spectral abundance, STEP achieved 80% precision at 5% recall, compared to MS2DeepScore’s 31.5% precision, for a more than 2-fold improvement. In contrast to spectrum-level evaluation, MS2DeepScore reached higher levels of precision, so we compared at a higher fixed precision of 60%. STEP achieved 10% recall, while MS2DeepScore achieved 2% recall. The overall structure-level AUPRC shows STEP at 25.3% versus MS2DeepScore’s 6.6%: a ∼4-fold improvement.

We additionally explored the upper bound performance of MS2DeepScore-like approaches. Because it is trained to predict 2048-bit Morgan fingerprint, we sought to understand the best case performance if the structures were known. In other words, the Tanimoto-similarity PR curve (Green line) in **Fig. 1** represents an upper bound for these two methods in both evaluation schemes. STEP is not constrained by this bound, and in most cases, it consistently performs better than the Tanimoto curve at moderate recall.

In low-recall regimes (around <5% recall), we observed unexpected drops in the precision of STEP (**Fig. 1**), given the assumption that model confidence is expected to increase monotonically as precision increases. We found that these unexpected drops in precision were often the result of incorrect prediction of structures with MT-GEM distances of 2 or 3 on regions of the molecule that were highly similar (21 out of 39 total cases). Such structural changes would be expected to result in nearly identical MS/MS if it were a single structural modification in the same region of the molecule. **Fig. 2** demonstrates two cases where multiple modifications occur near each other, producing minimal fragmentation differences, explaining precision drops in high-confidence regimes. These incorrect high-confidence predictions involving structures with MT-GEM distances of 2 or 3 accounted for more than half of the high-confidence predicted MT-GEM distance 1 MS/MS pairs in the testing (see **Fig. S1**). A broader case analysis of model behavior across combinations of STEP confidence and modified cosine similarity is provided in **Figs. S2-S7**.

**Fig. 2.**
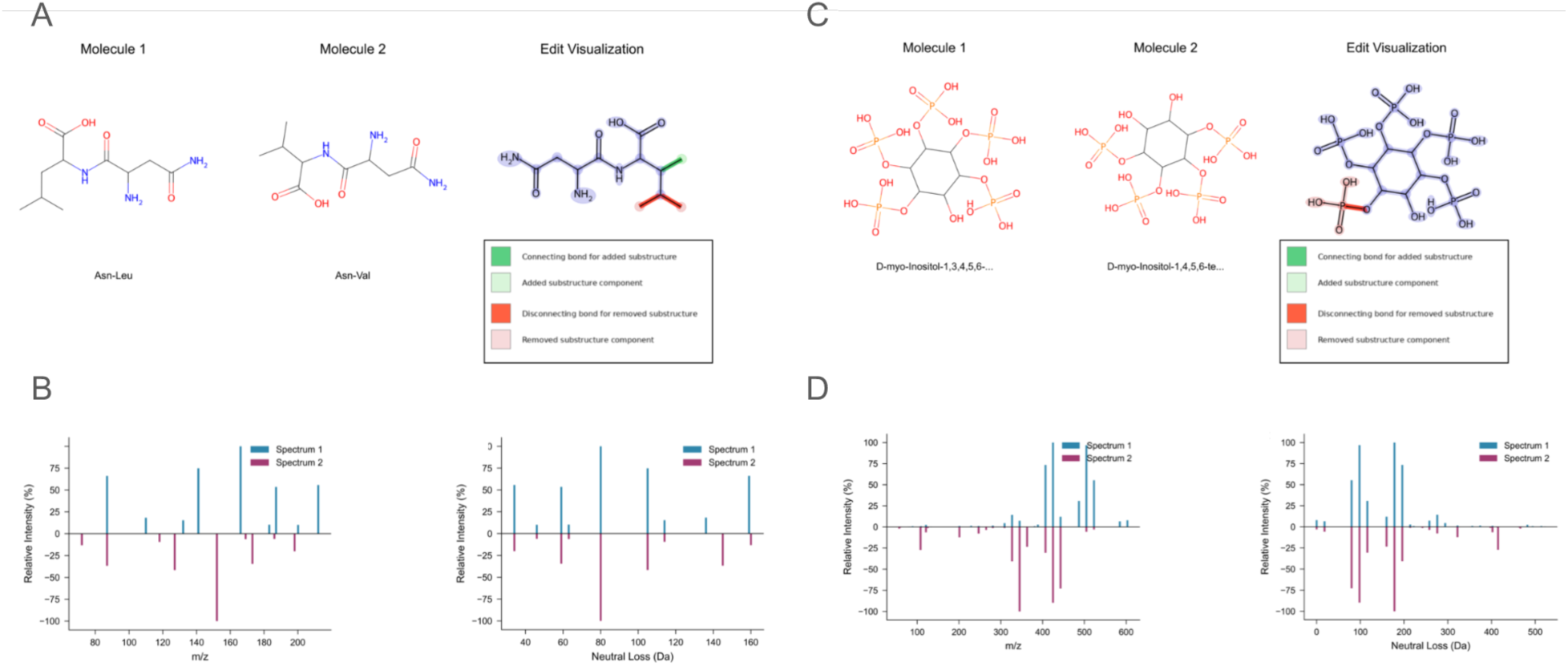
Comparison of MT-GEM predictions under high spectral similarity conditions, illustrating a false positive case (A–B) and a true positive case (C–D). Both pairs exhibit high STEP confidence (0.870) and high modified cosine (>0.93). (A) Molecular structures and edit visualization for a false positive pair (predicted MT-GEM = 1, ground truth = 3). The transformation requires two node removals (red) and one node addition (green), yielding a net change of −1 carbon that spectrally mimics a single edit operation. (B) MS/MS (upper) and neutral loss (lower) spectrum mirror plots for the false positive pair. High spectral similarity across both representations reflects the minimal net mass difference, obscuring the underlying three-edit transformation. (C) Molecular structures and edit visualization for a true positive pair (predicted MT-GEM = 1, ground truth = 1). The single substructure removal (red) represents an unambiguous graph edit directly reflected in spectral features. (D) MS/MS (upper) and neutral loss (lower) spectrum mirror plots for the true positive pair. Characteristic peak shifts correspond to the single structural modification, demonstrating expected model behavior when edits are not confounded by offsetting operations.

### Effect of prior probability and training coverage on model performance

We hypothesized that STEP performance varied as a function of precursor mass difference, because some precursor mass shifts associated with particular chemical transformations might be more common than others in experimental data. To assess whether STEP’s performance across precursor bins could be explained purely by differences in imbalanced data distribution or training coverage, we examined the relationship between prior probability, training set size, and average purity (**Fig. 3**). At the spectrum-level, grouping test pairs into 1.0 Da precursor mass-difference bins revealed no strong correlation (r=0.278) between the prior probability (fraction of MT-GEM distance 1 pairs) in each bin and the corresponding average precision on GNPSNIST-Test (**Fig. 3A**). Bins with similar prevalence can differ markedly in precision, and some bins with relatively low prior probability still achieve high precision. The same pattern was also observed when precision was averaged per unique structure pairs (**Fig. 3B**), indicating that these effects were not simply due to a few well-sampled structures. We next examined whether test performance was simply a function of how many examples the model had seen during training by plotting average precision versus the number of training spectrum pairs per mass-difference bin (1.0 Da) (**Fig. 3C**) and unique training structure pairs (**Fig. 3D**) per mass-difference bin (1.0 Da). Apart from bins with zero positive labels in the evaluation set, there was again no strong correlation (r = −0.092 and r = −0.084, respectively) between training set size and test performance. Some heavily sampled bins showed low precision, whereas others with relatively few examples performed well. These results show that neither the prior probability of MT-GEM distance 1 pairs nor the amount of training data fully explains the variation in STEP’s performance across precursor mass differences, indicating that the model actually captures additional structure-spectrum features beyond simple frequency effects.

**Fig. 3.**
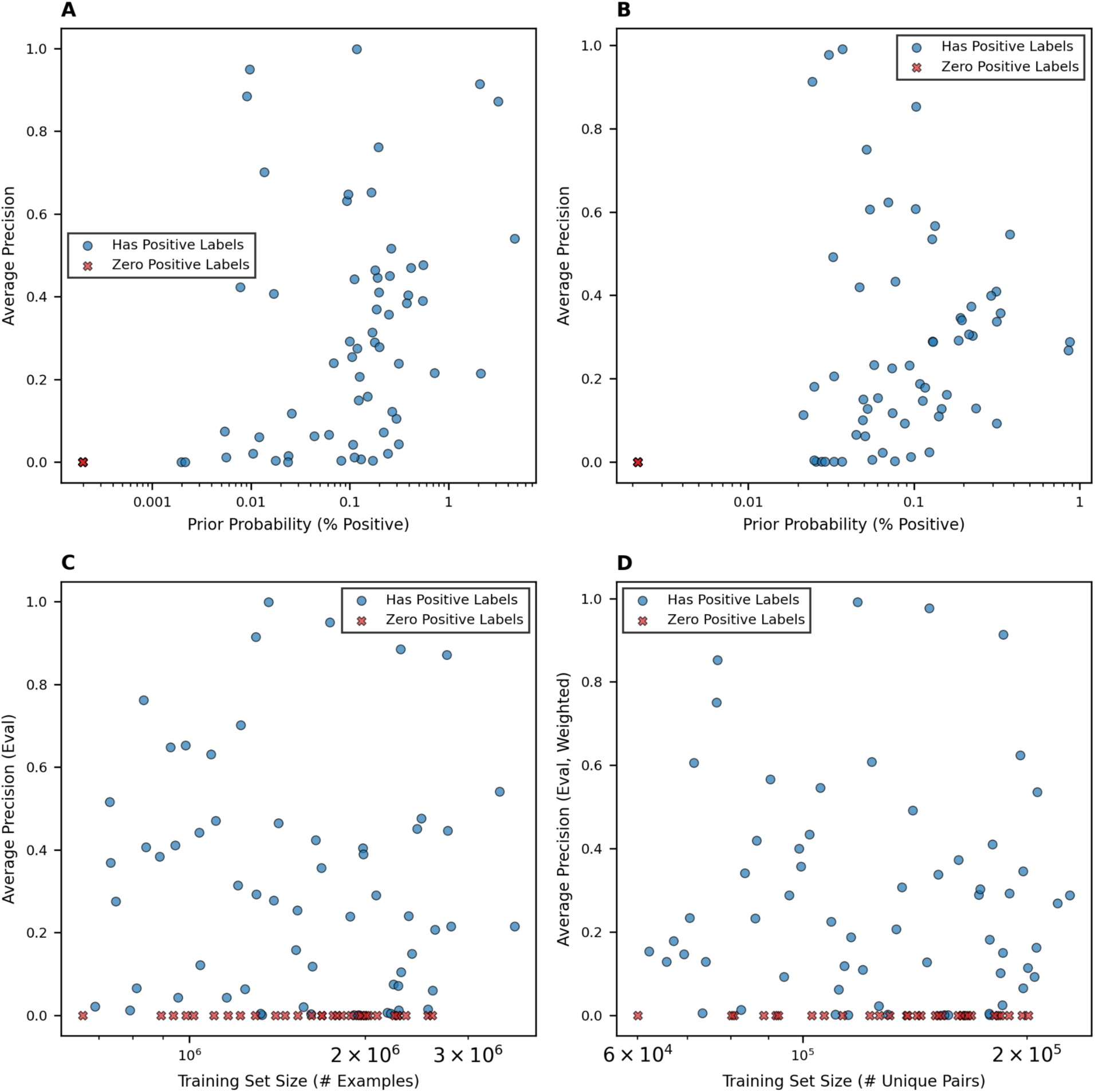
Positive label prevalence in GNPSNIST-Test (top, x-axis) and positive training examples prevalence in GNPSNIST-Train (bottom, x-axis) against the average precision of STEP on the precision recall curve for 1.0 Da width precursor mass difference bins on GNPSNIST-Test. Bins with no positive samples are labeled as red Xs while bins with positive samples are labeled as blue circles. (A) Scatterplot of prior probability of positive samples against average precision, log-scaled. (B) shows the same setup, but with unique structure pairs instead, to examine if the effect on prior probability appears on the structure pair level as well as the spectrum pair level. There is no strong correlation (r=0.061) between increased percent prevalence of positive samples on GNPSNIST-Test and model performance for both plots, likely indicating that STEP’s performance on precision recall plots is not most strongly determined by the prevalence of MT-GEM 1 pairs in data. (C) Scatterplot showing training set size (# of unique spectrum pairs) of MT-GEM 1 samples (x-axis) plotted against model performance on the test set. There is no strong correlation (r=-0.092) between the count of training samples and average precision. (D) Scatterplot showing training set size (# of unique structure pairs) of MT-GEM 1 samples (x-axis) plotted against model performance. There is no strong correlation (r=-0.084) between the count of training samples and average precision.

### Model performance under different instrument factors

To examine how different MS/MS conditions affect STEP, we evaluated model performance by instrument platform, adduct type, and compound class. In all analyses, precision and recall were averaged per unique structure pair to control for imbalances in the number of spectra per structure. We first separated test-set pairs by mass spectrometer platform to evaluate instrument-specific performance. QTOF acquisitions achieved the highest overall performance, with an average precision of 0.444 compared to 0.236 and 0.247 for Orbitrap and FTMS, respectively (**Fig. S8**). The QTOF precision–recall curve remained above the Orbitrap and FTMS curves across most of the recall range, indicating that MT-GEM distance 1 relationships could be detected more reliably using QTOF data. The higher performance observed for QTOF data suggested that differences in spectral quality and fragmentation characteristics between instruments may have an impact on STEP performance. Quadrupole-only data were excluded due to limited sample size.

Next, we grouped structure pairs by NPClassifier’s (26) pathway category to assess compound class performance. Carbohydrate and fatty-acid pairs showed the strongest performance, with average precision values of 0.690 and 0.486 and PR curves that maintained high precision at moderate recall (**Fig. S9**). Amino-acid/peptide and terpenoid pairs displayed intermediate performance, whereas alkaloids performed slightly worse. In contrast, shikimate and phenylpropanoid pairs exhibited poor performance (average precision 0.076). These results indicated that the recognizability of single-step structural transformations varies widely across biosynthetic families, likely reflecting differences in functional-group diversity and fragmentation regularity.

Finally, we evaluated performance as a function of adduct type by restricting to the two most prevalent species, [M+H]+ and [M+Na]+. Protonated pairs achieved substantially higher average precision (0.616) than sodiated pairs (0.438), with consistently higher precision at nearly all recall levels (**Fig. S10**). Because [M+H]+ accounted for approximately 85% of annotated adducts in GNPSNIST, this discrepancy is expected and likely reflects both greater training data coverage and differences in fragmentation chemistry between protonated and sodiated ions.

### Application to a synthetic gut community drug metabolism study

To evaluate STEP in a realistic experimental setting, we applied the model to a published functional metabolomics study (22) of drug metabolism by a defined human gut microbial community (MassIVE dataset MSV000094899). In this system, a 20-member synthetic community (Com20) comprising 20 commensal bacterial strains spanning six phyla and 17 genera, collectively encoding approximately 61% of the metabolic pathways found in a healthy human gut microbiome, was cultured under anaerobic conditions and challenged with a panel of clinically used small-molecule drugs. We evaluated STEP on this study dataset following the evaluation procedure described in Methods (see Methods **Evaluation on Experimental data**). Briefly, we constructed the Feature Based Molecular Network (FBMN) for the dataset and collected structural annotations from GNPS2 library matches. MT-GEM distances were computed between annotated structures and candidate spectrum pairs were generated using our standard pairing rules (see Methods **Evaluation on Experimental data**) without adduct filtering STEP scores for these pairs were then used to construct adduct-agnostic and adduct-aware precision–recall curves for predicting MT-GEM distance 1 relationships, while FBMN edges intersected with the same library-annotated pairs provided baseline performance in PR space. Overall, STEP achieved higher precision at matched recall than FBMN (STEP precision of 0.42 compared to FBMN precision of 0.25 at the same recall of 0.16) and maintained useful precision (>0.4) over a broader recall range (**Fig. S11**), indicating that directly modeling structural edit relationships from MS/MS spectra provides more specific connectivity than feature-level spectral similarity alone in this synthetic gut drug–metabolism setting.

Next, to illustrate STEP’s potential for practical structure elucidation, we used high-confidence (>0.7) MT-GEM distance-1 predictions between library-matched and previously unannotated spectra as candidates for microbially derived drug metabolites. Among the twelve drugs in the Com20 study, we focused on simvastatin as a representative example. The parent drug is confidently annotated based on MS/MS similarity at m/z 419.2792 (theoretical = m/z 419.2792), while an abundant feature at m/z 435.2743 had been proposed in the original work as a simvastatin-derived metabolite but lacked a definitive structural assignment. In the FBMN graph, these two features were associated with different network components (**Fig. 4A** nodes 12430 and 5538, respectively), meaning a cosine-based network did not suggest a direct relationship between them and lacking guidance for annotating the unknown feature. In contrast, STEP assigned the simvastatin pair (419.2792 and 435.2743) a high probability (0.87) of being an MT-GEM distance-1 relationship.

**Fig. 4.**
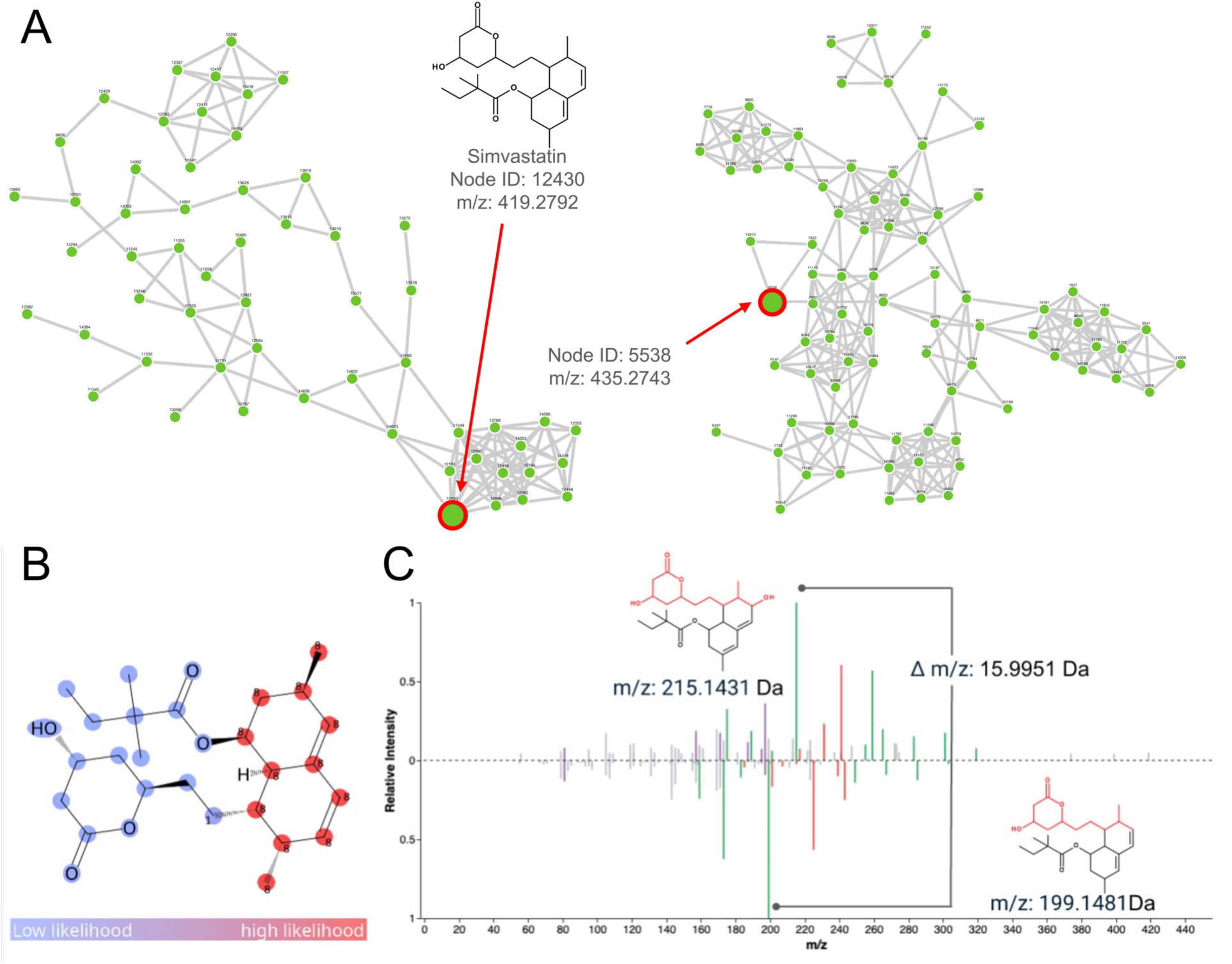
Edit-based annotation of a simvastatin-derived metabolite in the Com20 synthetic gut-community study. (A) Feature-based molecular networking (FBMN) view of the Com20 dataset. The node corresponding to simvastatin (m/z 419.2792, left red node) and the unknown feature at m/z 435.2743 (right red node) reside in different network components, so cosine-based networking does not suggest a direct relationship between them. (B) ModiFinder modification-site probabilities for the +15.9951 Da edit between simvastatin and the unknown feature. Colors encode per-atom likelihood (blue, low; red, high), highlighting the distal aromatic/statin core ring as the most probable site of mono-oxygenation and assigning low probability to the lactone side chain. (C) Fragment-alignment mirror plot with MAGMa annotation for the unknown (upper) and known (lower) spectra. The unknown spectrum is annotated using the predicted hydroxylated simvastatin structure. Bars are colored by fragment class: unaligned peaks (grey), unannotated shifted peaks (red), annotated shifted peaks (green), and annotated unshifted peaks (purple). The most intense shifted peak pair (m/z 215.14, unknown; m/z 199.15, known; Δm/z = 15.9951 Da) corresponds to fragments containing the distal aromatic ring, with fragment-structure highlighting (insets) confirming that the cleavage site is adjacent to the predicted hydroxylation position.

The precursor mass difference of 15.9951 Da was consistent with the addition of a single oxygen atom, indicating that the unknown feature is likely a monohydroxylated simvastatin analogue rather than a multi-step degradation product. We then integrated ModiFinder (19) to localize this putative +O modification on the simvastatin scaffold. ModiFinder enumerated single-step modifications compatible with the observed mass shift and returns per-atom likelihoods for the edit location. For this pair, it highlighted the distal aromatic ring of the statin core as the highest-probability region for hydroxylation, while assigning low probability to the lactone side chain. This prediction matched the manual annotation proposed by expert chemists who concluded that the unknown corresponds to a hydroxylated simvastatin with an OH group added to the terminal ring (22). Combinatorial fragmentation further supported this interpretation by annotating the highest-intensity shifted peak pair between simvastatin (m/z 199.15) and hydroxylated simvastatin (m/z 215.14) as corresponding to the same substructure, differing only by a single oxygen addition (**Fig. 4C**). More specifically, these fragments corresponded to substructures containing the distal aromatic ring, indicating that the cleavage producing these fragments occured adjacent to the predicted hydroxylation site. Fragments reporting on the lactone moiety remain largely unshifted. This fragment-level evidence was consistent with the ModiFinder prediction (**Fig. 4B**), providing orthogonal confirmation that the +O modification occurs on the aromatic ring rather than elsewhere on the molecule.

This case study demonstrated that STEP can recover a one-step structural relationship between a known drug and an unannotated feature that is invisible to FBMN, and that combining STEP with ModiFinder and fragment-level analysis rapidly narrows this relationship to a concrete structural hypothesis. Whereas the original study required a multi-step workflow involving additional experimental work, network inspection, and extensive manual MS/MS interpretation to propose this metabolite, our computational framework surfaced the candidate pair automatically, suggested the specific type of modification, and localized its most likely site, substantially shortening and simplifying the path from unknown peak to structural annotation.

### Application to bioactivity-guided natural product analogue mining

As a natural products case study, we applied STEP to a bioactivity-guided LC–MS/MS dataset (23) acquired from solid-phase extraction (SPE) fractions of extracellular metabolites produced by *Serratia marcescens* DSM 1636 and DSM 30121 (See Methods **Acquisition of the Serratia Marcescens Dataset**). Fractions were screened for antagonistic activity against oomycete phytopathogens and profiled by short-gradient LC–MS/MS in positive ion mode. MS/MS data were processed in MZmine and used to construct a GNPS2 feature-based molecular network (FBMN), onto which bioactivity was mapped to prioritize activity-associated molecular families.

To explore how STEP facilitated analogue prioritization within bioactivity-associated molecular families, we examined high-confidence (>0.7) MT-GEM distance-1 predictions centered on a feature annotated in FBMN as serratiochelin (m/z 430.1610, **Fig. S12**). Within the same component, STEP identified multiple candidate one-step analogues, e.g., Node ID 20096 (m/z 444.1764), which, although located within the same component in FBMN, was not directly connected to the serratiochelin anchor in the molecular network. The ion m/z 444.1764 showed a precursor mass difference of 14.0147 Da relative to the anchor (consistent with a single CH2 increment). This MS/MS pair exhibited a low modified cosine score of 0.5060, insufficient to generate a direct edge in the molecular network, but produced an MT-GEM distance-1 likelihood of 0.8. For this pair, downstream modification analysis using ModiFinder suggested a putative N-methylation of the central amine as the most plausible single-step edit (**Fig. 5 A and B**).

**Fig. 5.**
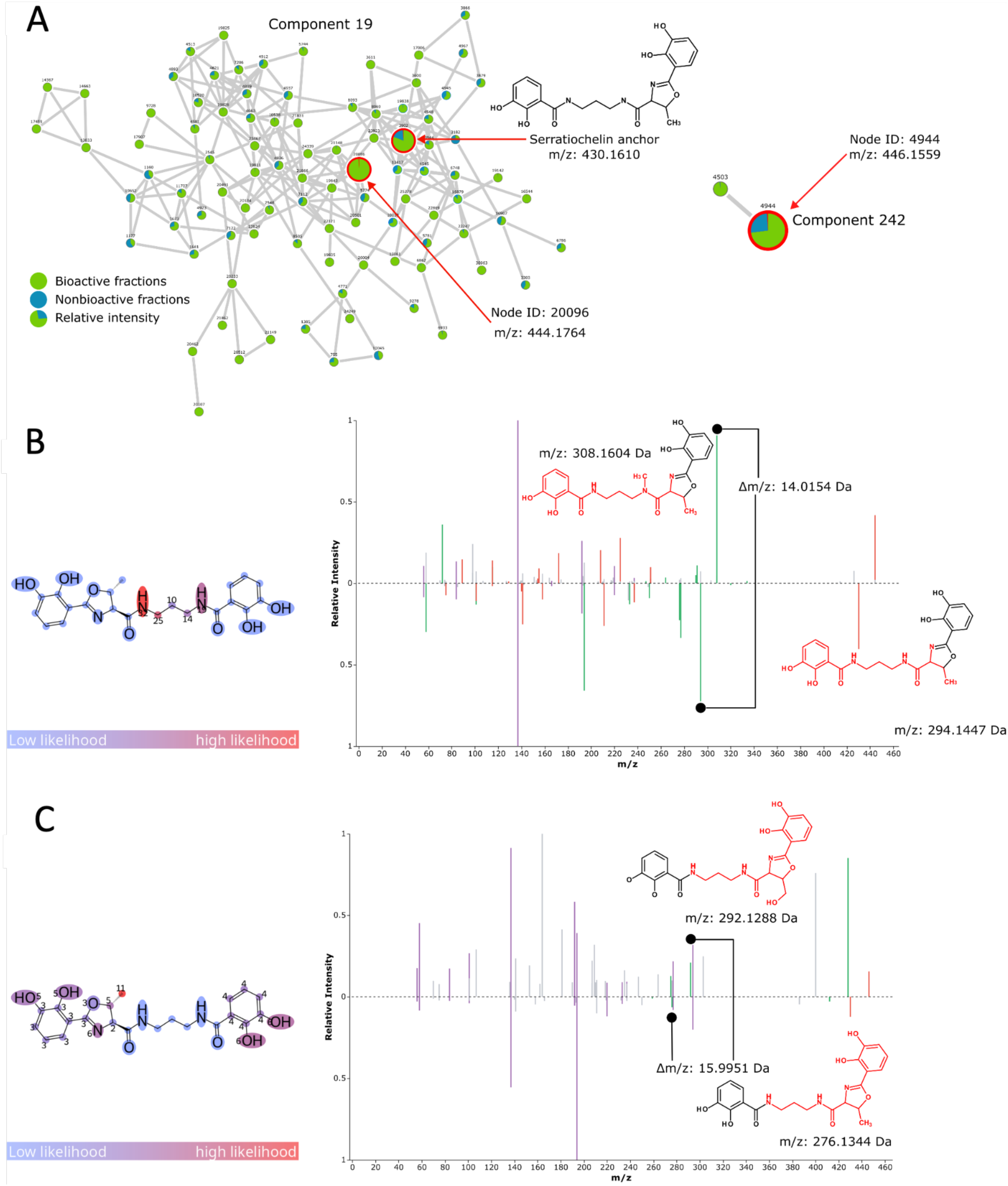
Edit-based analogue prioritization within a bioactivity-guided *Serratia marcescens* metabolomics dataset. (A) Feature-based molecular network (FBMN) of SPE fractions from *S. marcescens* DSM 1636 and DSM 30121, with bioactivity mapped onto nodes. Node color indicates fraction type (green, bioactive; blue, nonbioactive) and the pie chart within each node reflects the relative ion intensity across fractions. Two network components are shown: Component 19, containing the serratiochelin anchor (m/z 430.1610) and Node ID 20096 (m/z 444.1764), a candidate distance-1 analogue not directly connected to the anchor in the molecular network; and Component 242, containing Node ID 4944 (m/z 446.1559). (B) Mirror plot and ModiFinder modification-site probabilities for the serratiochelin anchor – Node ID 20096 pair (Δm/z = 14.0154 Da). Colors encode per-atom likelihood (blue, low; red, high), with the central amine highlighted as the most probable site of methylation, consistent with a single CH2 increment. (C) Mirror plot and ModiFinder modification-site probabilities for the serratiochelin anchor – Node ID 4944 pair (Δm/z = 15.9951 Da), with the C5 methyl group of the oxazoline ring identified as the most probable site of hydroxylation, consistent with the addition of a single oxygen atom.

In contrast to the within-family analogues, STEP predicted a distance-1 relationship between the serratiochelin anchor and Node 4944 (m/z 446.1559), which was not grouped within the same component but was predominantly detected in bioactive fractions, suggesting a potential contribution to the observed biological activity (**Fig. 5A**). Despite the absence of cosine-based connectivity (cosine similarity = 0.4264), STEP assigned a strong single-edit likelihood to this pair (Probability = 0.870). The 15.9951 Da precursor mass shift relative to the anchor was consistent with the addition of a single oxygen atom. Downstream modification analysis using ModiFinder localized this change to the methyl carbon at C5 of the oxazoline ring, supporting a putative hydroxylation of the threonine-derived methyl group (**Fig. 5C**). This modification may arise from post-cyclization oxidation, differential substrate incorporation during biosynthesis, or a non-canonical transformation during oxazoline ring formation (27).

## Methods

### Data Acquisition and Preprocessing

We trained a binary classifier to predict whether pairs of MS/MS spectra correspond to molecules separated by a single MT-GEM transformation. Training required generating labeled pairs of spectra with known MT-GEM distances calculated from annotated molecular structures. We compiled tandem mass spectra from the GNPS2 spectral database and the National Institute of Standards and Technology 2020 Mass Spectral Library (NIST20) (20). We refer to the union of these datasets as GNPSNIST. We used matchms to filter spectra to retain only positive mode small molecule data in the mass range 10-1000 m/z, and removed peaks below 0.1% intensity as noise, filtered for a minimum of 5 peaks to ensure sufficient structural information. We truncated spectra to the top 100 most abundant peaks to focus on structurally relevant information. After the initial filtering, 490,292 spectra resulted. Following preprocessing steps from Bui et al.’s TransExION (11) implementation, we applied a 3.0 m/z sliding window to remove dense peak clusters, normalized peaks from 0.0 to 1.0 relative to maximum intensity, and retained float32 precision for all measurements to preserve subtle mass differences. Prior to dataset splitting, we retained 416,099 mass spectra representing 31,344 unique structures determined by the first 14 characters of each InChiKey.

To prevent data leakage and model memorization of substructure information during training, we employed the dataset splitting methodology validated by Strobel et al. (28). This approach partitioned data by maximizing pairwise Tanimoto distance between molecular structures across train and test sets, calculated using 2048-bit RDKit Daylight fingerprints. Train-test similarity for each test structure was defined as the maximum pairwise similarity between that structure and all training structures. The splitting algorithm proceeded in three steps: (1) selecting structures with low train-test similarity by binning all structures into 13 equally spaced train-test similarity bins from 0.4 to 1.0 and randomly sampling structures from each bin, (2) selecting structures with high pairwise similarity by constructing a graph where edges connect structures with Tanimoto similarity greater than 0.7, then performing random walks from high-degree nodes to ensure coverage of similar structure pairs in the test set, and (3) iteratively removing structures from the training set that exceed the similarity threshold for each bin to enforce low train-test similarity across bins. This process continued until all bins reach their target allocation or 80% of the training set is removed. The resulting split ensures the test set contains structures at varying distances from the training set while maintaining representation of structurally similar pairs for evaluation.

To ensure consistent fragmentation patterns between spectral pairs, we required exact matching of instrument type, ionization type, and adduct, and constrained collision energy differences to +-5eV maximum. We found that 98% of single MT-GEM distance pairs were constrained to the 14-104 delta precursor m/z range, so the training data was constrained to this range on the assumption that it was representative of the true distribution of delta mass differences. These filtering criteria ensured that high-quality spectra are used for model training and that high quality spectrum pairs with comparable fragmentation conditions are used for model training. After pair generation, we observed severe class imbalance with <0.5% of pairs representing single structural modifications (MT-GEM distance of 1), which was expected given the diverse, often unrelated molecular structures in the dataset.

### Evaluation Protocol for GNPSNIST

For STEP, we used the predicted probability from the sigmoid activation as the confidence score, without additional calibration. For each baseline method (modified cosine, spectral entropy similarity, MS2DeepScore, and DreaMS), we computed pairwise similarity scores for all test pairs and used these scores as confidence thresholds for predicting MT-GEM distance of 1. As a structural reference baseline, we additionally computed 2048-bit Morgan fingerprint (radius 2) Tanimoto similarity from the known molecular structures of both spectra. Because this metric requires structural knowledge unavailable for unannotated features, it represents an upper bound on the performance achievable by methods trained to predict fingerprint similarity. Precision-recall curves were generated by sweeping across all unique confidence thresholds for each method.

We reported performance under two evaluation schemes. Under spectrum-level evaluation, each of the 3,880,795 spectral pairs contributed independently to the precision-recall calculation. Because some structure pairs were represented by many replicate spectra, a single structure pair with n and m spectra generates up to n × m spectral pairs, allowing high-multiplicity pairs to disproportionately influence spectrum-level metrics. We reported the area under the precision-recall curve (AUPRC), computed via trapezoidal integration of the precision-recall curve, as the primary summary metric for this scheme.

To control for this uneven spectral representation effect, we additionally report structure-level evaluation. For each unique structure pair, defined by the sorted canonical SMILES pair, we assigned a weight inversely proportional to the number of spectral pairs representing that structure pair. AUPRC was then computed with these sample weights, ensuring that each structural relationship contributes equally regardless of spectral multiplicity.

### MT-GEM Distance Calculation

The Molecular Transformation Graph Edit Measure (MT-GEM) quantifies structural similarity through graph operations. For each molecular pair, we computed the maximum common substructure (MCS) using RDKit with computational constraints: limiting calculations to molecules with fewer than 60 heavy atoms, requiring minimum Tanimoto similarity of 0.1 to avoid expensive operations on unrelated structures, and requiring the MCS to contain at least half the atoms of both structures. The MT-GEM distance equals the sum of minimum edge removals required to isolate the MCS from remaining substructures in each molecule, providing a direct proxy for chemical transformations where MT-GEM distance of 1 indicates a single substructure modification. **Fig. 6** illustrates the MT-GEM calculation through representative molecular transformations, showing how substructure removals quantify discrete chemical modifications.

**Fig. 6.**
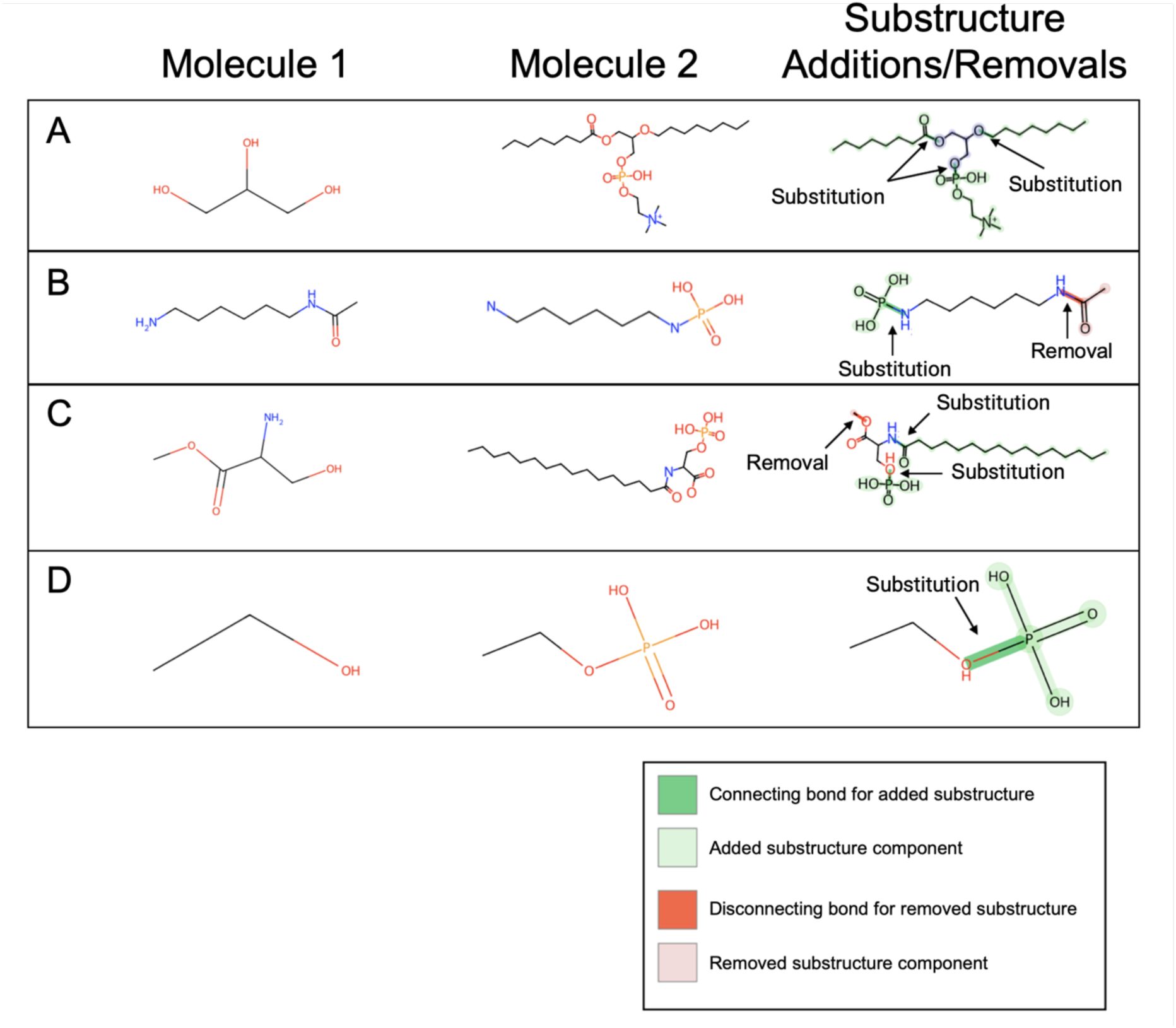
Illustration of MT-GEM distance through representative molecular transformations. The substructure “Addition” and “removal” refer to graph-level substructure operations (attachment or detachment of connected components at a single bond for heavy atoms with H omitted in the graph). We acknowledge that graph-level additions here are in chemical terms substitutions, as the added substructure replaces an implicit hydrogen. (A) Glycerol to phosphatidylcholine (MT-GEM = 3): sequential addition of two fatty acid chains and phosphocholine headgroup. (B) N-acetyl-hexanediamine to N-phospho-hexanediamine (MT-GEM = 2): acetyl removal from one terminus, phosphate addition to the other. (C) Serine methyl ester to N-palmitoyl-O-phosphoserine (MT-GEM = 3): methyl ester hydrolysis, N-palmitoylation, and O-phosphorylation. (D) Ethanol to ethyl phosphate (MT-GEM = 1): phosphate addition.

### Evaluation on Experimental data

We evaluated our model on experimental data by validation against feature-based molecular networking performance as baselines. To establish baseline performance, we used spectra library-matched via GNPS2 to previously identified structures with SMILES annotations. From this set of SMILES, we calculated the pairwise MT-GEM distance between all structures to use as an evaluation set per dataset. We paired all spectra based on our established criteria (See Methods **Data Acquisition and Preprocessing**), but without adduct filtering to enable de novo analysis without requiring expert annotation for inference. From this set of paired spectra, we mapped the known MT-GEM distances to library-matched spectra to create a sub-dataset of spectra identified through library matching. We ran inference with our best performing STEP model based on validation loss, then plotted the precision-recall curve for non-adduct filtered vs adduct filtered to see the hypothetical performance difference between if we know the adducts in unknown spectra compared to if we do not. We also plotted the molecular networking precision and recall as a single point on the figure by intersecting pairs of structures that exist in the Feature-Based Molecular Networking (FBMN) output based on library-matched spectra, which we consider known spectra. If a structure pair existed in the FBMN output pairs, it was considered a “prediction” for the sake of the precision-recall chart.

### Dataset Processing and Threshold Selection on Experimental Data

For each validation dataset, we applied consistent preprocessing and evaluation criteria. Feature-Based Molecular Networking (FBMN) datasets underwent pairwise spectral matching of all output spectra. All spectral pairs were filtered to match our training criteria: identical collision energy (±5 eV), instrument, and ionization mode. FBMN datasets proceeded without adduct filtering, as this information is unavailable in feature-based workflows.

Model inference on all qualified pairs used thresholds selected through precision-recall analysis against library-matched spectra containing annotated SMILES structures. This validation approach provides dataset-specific performance baselines, enabling chemists to calibrate confidence in predictions.

### Deep Neural Network Architecture

We retrofit two state-of-the-art spectrum embedding architectures, Bui et al’s TransExION architecture (11) and Bushuiev et al’s DreaMS architecture (18), to predict single edit distance edges. At a high level, TransExION encodes hypothetical peak shifts into a matrix representation of the whole number and decimal place mass differences between peaks, allowing the model to attend to the fragmentation patterns of the two spectra, e.g., if 16.00 Da can be observed as a mass difference indicating a minor substructure modification of oxygen. We also utilize the DreaMS model pretrained on millions of unannotated MS/MS via predicting masked spectral peaks and chromatographic retention orders to generate rich embeddings for each spectrum (18). We concatenate the spectrum representations output by DreaMS concatenated with the larger of the two molecules first, then the smaller molecule’s embedding, then the result of subtracting the larger molecule’s embedding from the smaller molecule’s embedding to encode the structure difference between both spectra. We then feed the TransExION representation of the spectrum peak shifts and the concatenated embeddings into a transformer decoder with a CLS token placed at the beginning and separator tokens between the TransExION spectrum embeddings, TransExION neutral loss embeddings, and DreaMS vector conditioned on a learned linear projection of the precursor mass difference between the two spectra. The output of the transformer decoder is then fed into a linear prediction head that outputs the binary prediction logits for if the spectrum pair is a single edit distance apart. This process is illustrated in **Fig. S13**.

### Training Procedures

We trained our model, STEP, with the AdamW optimizer (29) with learning rate 2.5e-5, betas (0.9, 0.999), weight decay 1e-3, and epsilon 1e-7. We implemented cosine annealing with one cycle over a full epoch. To address severe class imbalance, we combined three strategies: focal loss to penalize confident misclassifications, artificial upsampling ensuring 2% of each batch contained positive examples to prevent gradient collapse, and label smoothing to mitigate overconfidence (30).

Training utilized dual A100 GPUs with a batch size of 512, requiring approximately 48 hours per epoch over 230 million training pairs. We trained for multiple epochs with early stopping based on validation set performance to prevent overfitting while ensuring the model captured generalizable transformation patterns.

### Acquisition of the Serratia Marcescens Dataset

Extracellular metabolites from Serratia marcescens DSM 1636 and DSM 30121 were obtained from liquid cultures grown in minimal medium at 27 °C and 150 rpm for 48 h. Culture supernatants were extracted with ethyl acetate (1:1, v/v). The organic phase was evaporated, and the resulting crude extract was fractionated by solid-phase extraction (Strata-XL polymeric reversed-phase, Phenomenex) using a stepwise methanol gradient (20–100%). Fractions were concentrated and used for downstream LC–MS/MS analysis.

Untargeted metabolomics data were acquired using a Vanquish UHPLC system coupled to a Q Exactive HF quadrupole-Orbitrap mass spectrometer (Thermo Fisher Scientific, Bremen, Germany) equipped with a heated electrospray ionization (HESI) source operating in positive ion mode. Chromatographic separation was performed on a Kinetex EVO C18 column (50 × 2.1 mm, 1.7 µm, 100 Å; Phenomenex) at 25 °C with a flow rate of 0.5 mL min⁻¹. Mobile phases consisted of water with 0.1% formic acid (A) and acetonitrile with 0.1% formic acid (B). The gradient started at 5% B, increased linearly to 50% B at 8 min and to 99% B at 10 min, held until 13 min, and returned to initial conditions for re-equilibration.

Data-dependent acquisition (DDA) was performed over an m/z range of 150–1500 with full MS scans at 30,000 resolution, followed by MS/MS scans at 15,000 resolution on the top 5 most intense precursor ions. Fragmentation was carried out using stepped normalized collision energies of 20, 25, and 30. Source parameters were set to 3.5 kV spray voltage, 250 °C capillary temperature, 400 °C probe heater temperature, sheath gas 50, auxiliary gas 12, sweep gas 1, and S-lens RF level 50.

## Discussion and Conclusion

In this study, we propose that reframing spectral similarity as a predictor of discrete structural edits changes how MS/MS data can be used. MT-GEM provides a physically motivated notion of distance based on substructure removals and additions, and STEP learns to recognize MT-GEM distance 1 relationships directly from pairs of MS/MS spectra. In our testing regime, STEP yields large gains over modified cosine, spectral entropy similarity, and Tanimoto score-based machine learning approaches such as MS2DeepScore, at both the spectrum and structure level, and in some cases even exceeds the performance of the oracle fingerprint Tanimoto curve. Analysis of prior probability and training coverage shows that these gains are not simply due to imbalanced training or testing data abundance. For example, precursor mass bins with similar prevalence and a similar number of training examples vary widely in precision, with some low-prevalence bins still perform well, suggesting that the STEP model actually learned additional structure-spectrum relations beyond simply being affected by training or testing data frequency.

The key innovation lies not in incremental performance improvements but in fundamentally changing how metabolite annotation proceeds. By providing discrete, interpretable predictions of structural transformations, we enable hypothesis-driven structure elucidation with a significantly reduced search space where chemists receive specific modifications to evaluate rather than abstract similarity scores to interpret. Combined with modification site prediction through tools like ModiFinder, this transforms a previously ad hoc problem into a systematic annotation process that provides increased annotation throughput. In the synthetic gut-community and bioactivity-guided natural product case studies, instead of manually exploring a large molecular-network neighbourhood and informally ranking many potential candidates, the expert chemist is presented with a small, prioritized set of “single-step” hypotheses linking known scaffolds to unknown features, together with suggested modification sites and diagnostic fragments to inspect. These examples illustrate how edit probabilities can support a more streamlined, high-throughput style of annotation, turning generic spectral relatedness into specific, testable chemical stories in a realistic microbiome–drug metabolism setting.

While we saw a significant improvement in STEP’s ability to predict MT-GEM, other alternative approaches, one key weakness is that insufficient negative mode spectrum data exists to sufficiently train the model. Therefore, STEP is limited to only handling positive mode MS/MS spectra. Additionally, STEP cannot distinguish stereoisomers, as these can produce identical fragmentation patterns despite what is inherited from tandem mass spectrometry. The present model also predicts only whether a pair has an MT-GEM-distance of one, without explicitly reasoning over full edit distances or ordered reaction sequences. Future work may address these aspects by extending the training data, developing models that predict richer edit information (for example, approximate MT-GEM distance, or likely sequences of edits), and incorporating additional experimental dimensions such as retention time, ion-mobility separation, and collision-energy dependence. This would allow the same edit-based framework to describe more complex biotransformations and provide more generalizable guidance. In parallel, there is a clear opportunity to integrate STEP more tightly with complementary tools such as ModiFinder, so that edit prediction, modification-site localization, and fragment-level validation are bundled into highly automated, high-throughput workflows. In such pipelines, edit-based scores would serve as a front end that routes unknowns to appropriate downstream modules, helping large metabolomics studies propagate structural hypotheses through molecular networks and ultimately accelerating community-wide efforts in metabolite annotation.

## Data Availability / Source Code

The code used in this manuscript is available at https://github.com/Wang-Bioinformatics-Lab/edit-distance-paper. The GNPS dataset is available at https://doi.org/10.5281/zenodo.11193897, and a version of the model weights trained only on the GNPS library spectra is made available at https://zenodo.org/records/19445015. The latest NIST dataset is available for purchase at https://www.nist.gov/programs-projects/nist23-updates-nist-tandem-and-electron-ionization-spectral-libraries.

## Acknowledgments

MW was supported by NIH grant 5U24DK133658. XW was supported by a UCR Fellowship. VVP was supported by NIH grant R35GM158024.

