## Supplementary material for "Predicting Discrete Structural Transformations in Small Molecules from Tandem Mass Spectrometry": Fig. S1, Fig. S2, Fig. S3, Fig. S4, Fig. S5, Fig. S6, Fig. S7, Fig. S8, Fig. S9, Fig. S10, Fig. S11, Fig. S12, Fig. S13, Table S1

**Affiliations:**

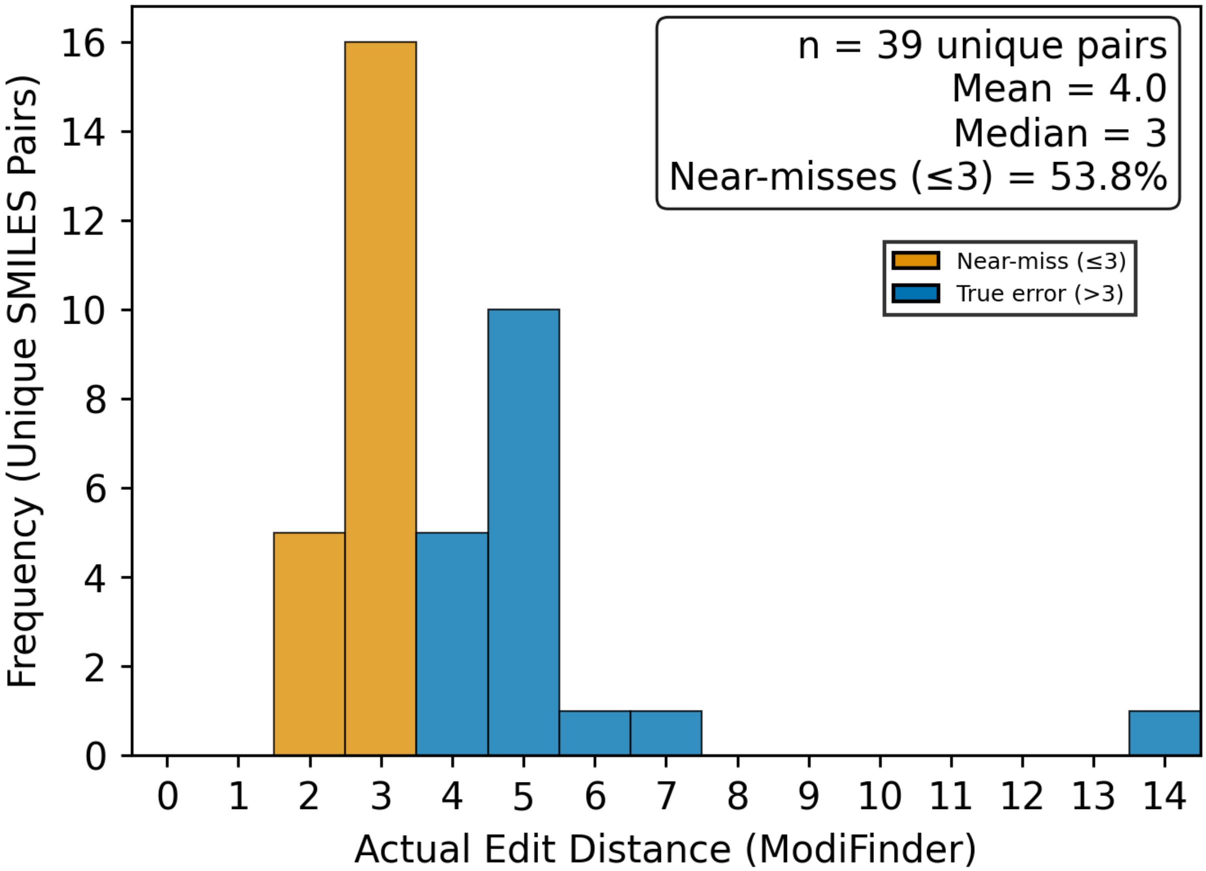


**Fig. S1.** Distribution of all false positive cases of true MT-GEM edit distances among high-confidence STEP distance-1 predictions. Histogram of actual edit distances (x-axis, computed with ModiFinder) for 39 unique SMILES pairs that STEP predicted as MT-GEM distance 1 with high confidence but were labeled as non-distance-1 in the GNPSNIST test set (False positive cases). Bars in orange denote “near-miss” cases with true edit distance ≤3, while blue bars indicate larger discrepancies (>3). Over half of the errors (53.8%) fall into the near-miss category, with many pairs having edit distances of 2–3, consistent with structural changes localized to similar regions of the molecule and expected to yield very similar MS/MS spectra.

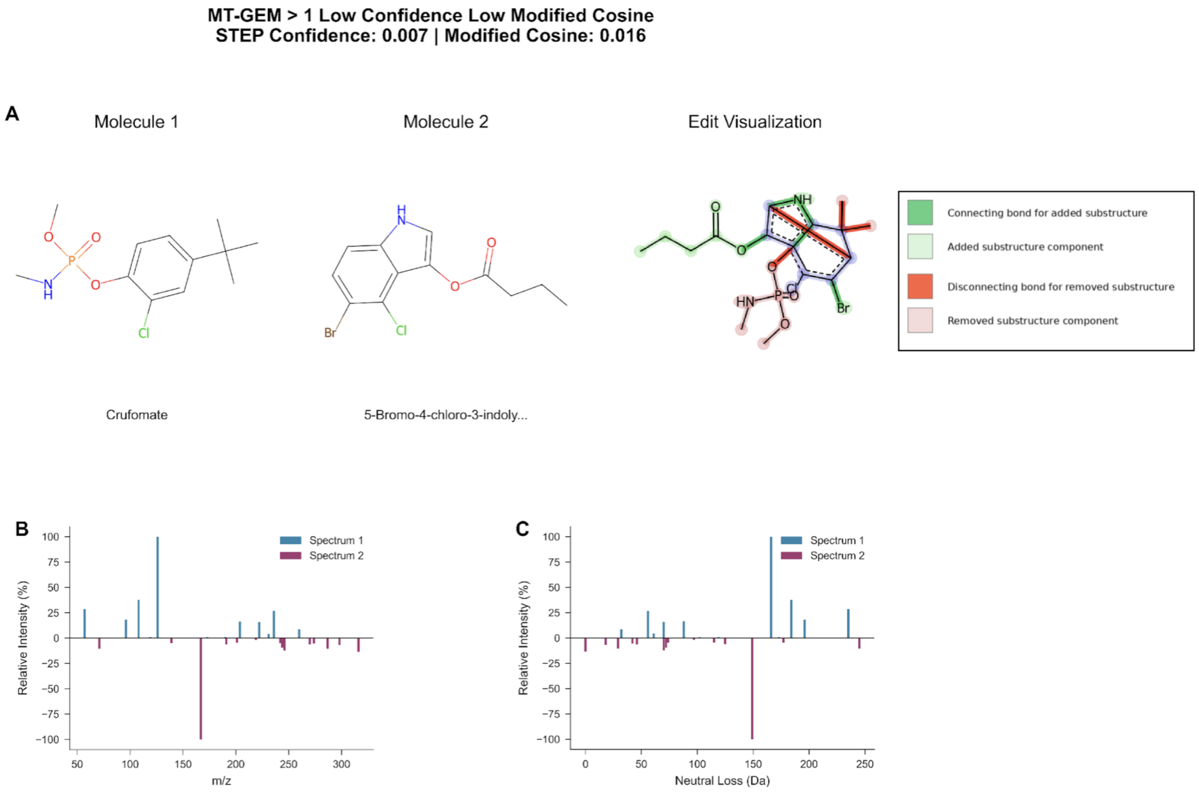


**Fig. S2.** This figure illustrates a low confidence negative example (MT-GEM > 1) where two unrelated molecules are correctly determined by STEP to be not a single edit apart, which is similarly reflected in the low modified cosine of 0.016.

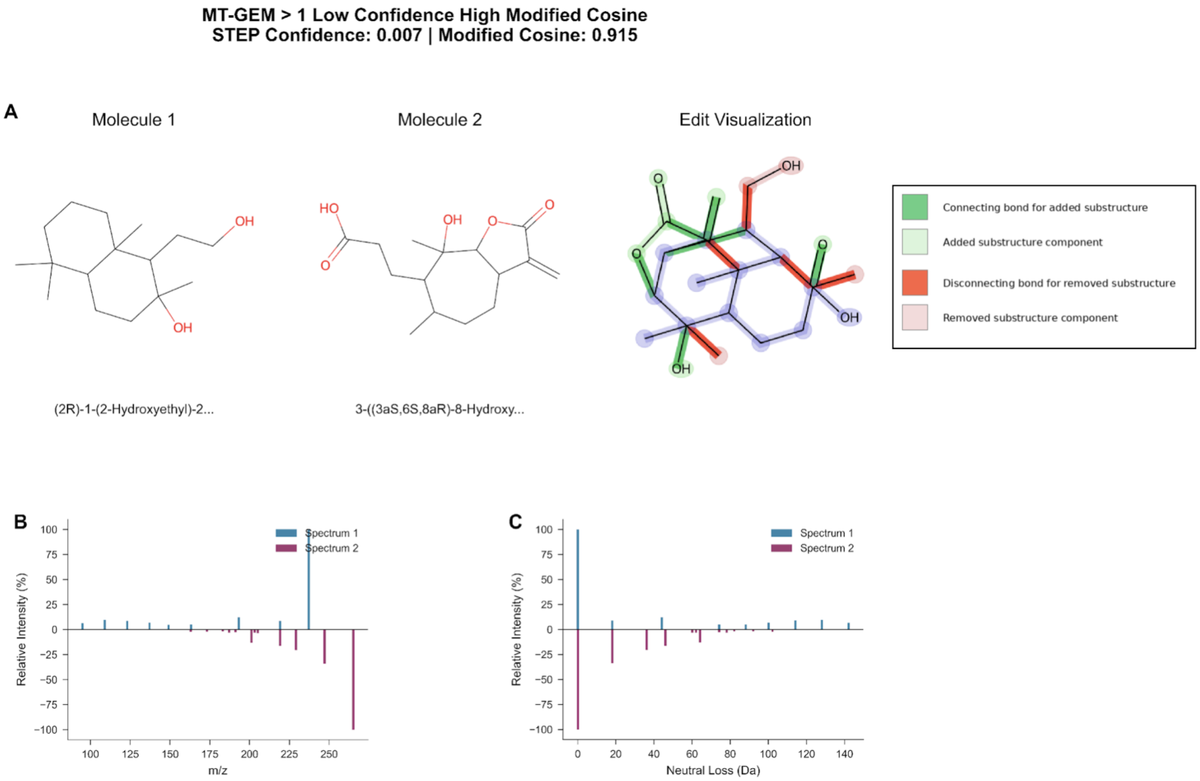


**Fig. S3.** This figure illustrates a low confidence negative example (MT-GEM > 1) where two unrelated molecules are correctly determined by STEP to be not a single edit apart, despite the fact that the spectrum pair implies a very high spectral similarity through a modified cosine of 0.915.

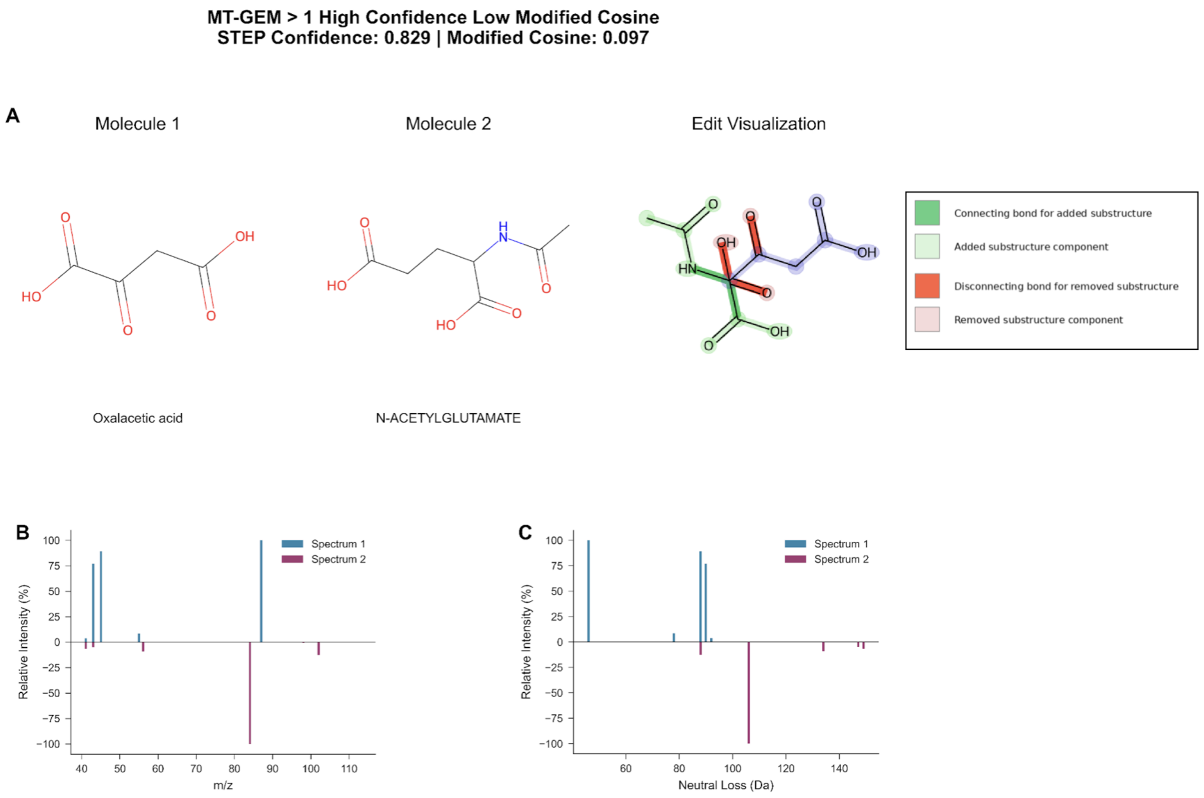


**Fig. S4.**. This figure illustrates a high confidence negative example (MT-GEM > 1) where two unrelated molecules are incorrectly predicted by STEP. The likely reason for this is that the model identified peak shifts characteristic of a single modification, specifically a peak shift matching the precursor mass difference, giving a high false positive signal.

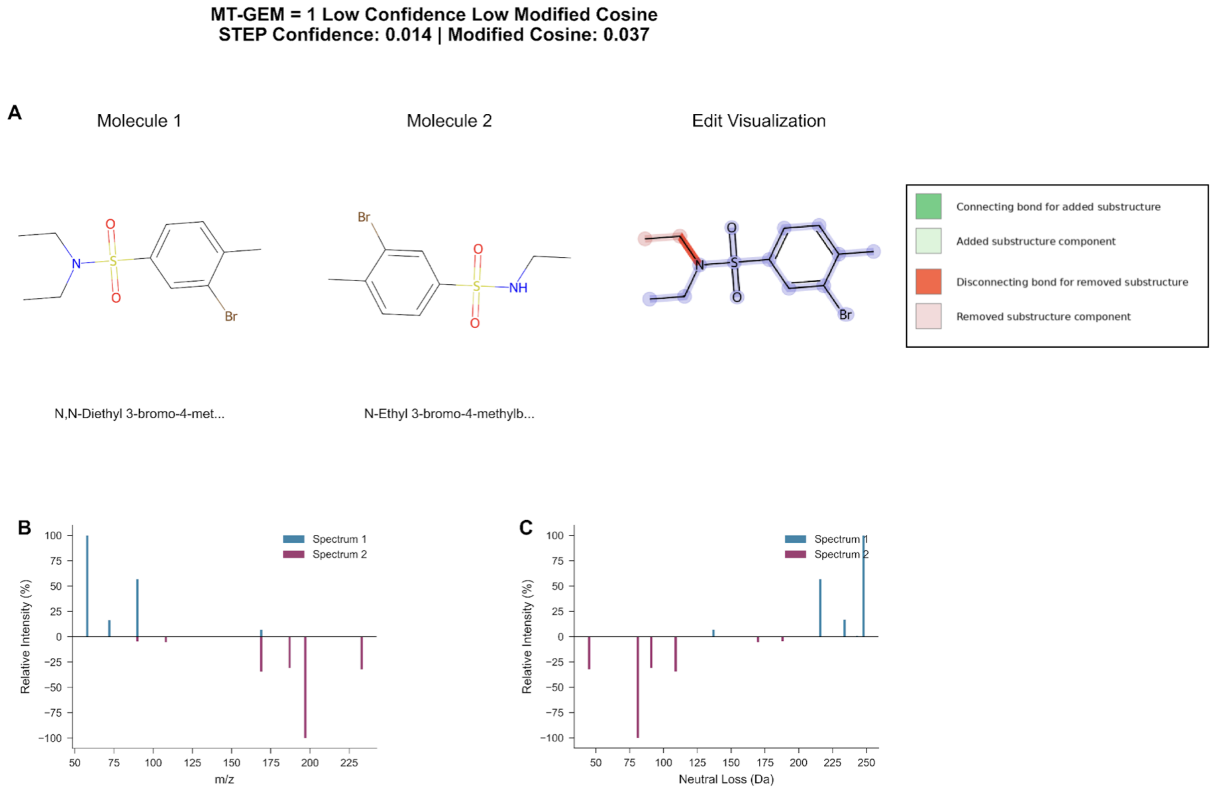


**Fig. S5.** This figure illustrates a low confidence positive example (MT-GEM = 1) where two related molecules are incorrectly predicted by STEP. The likely reason for this is that the model failed to identify peak shifts characteristic of a single modification, giving little to no signal to predict MT-GEM 1.

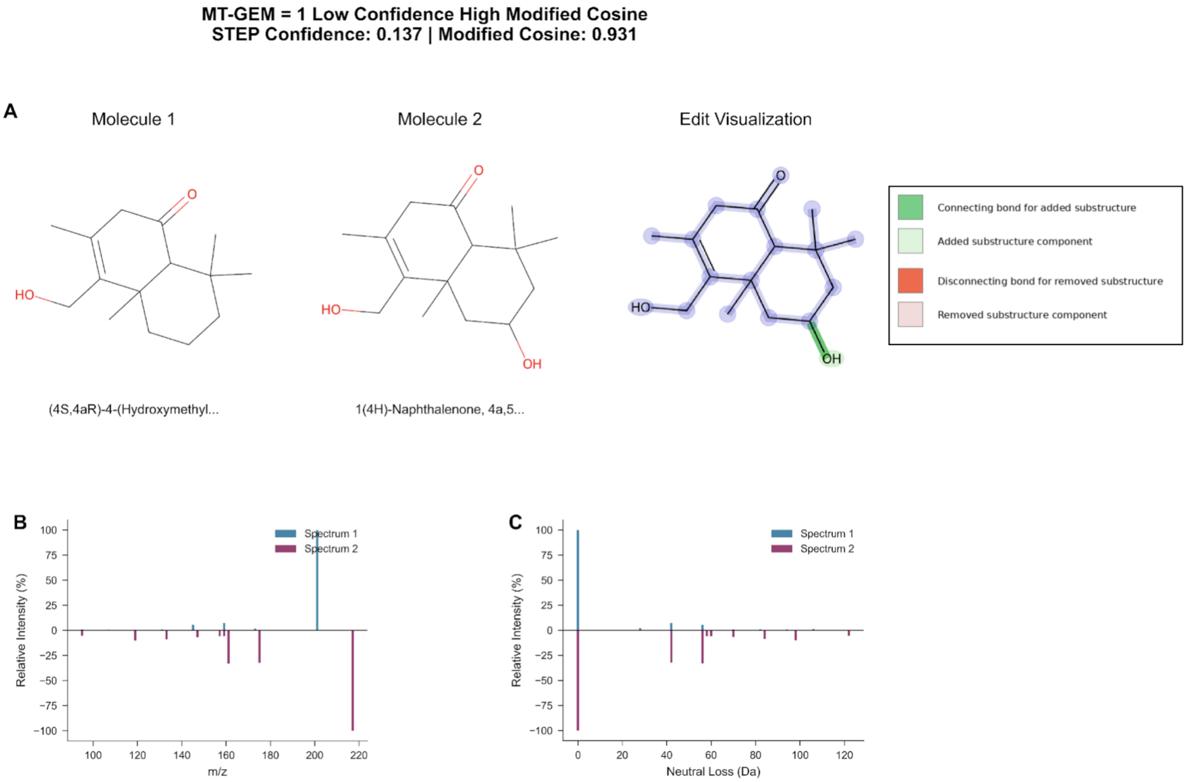


**Fig. S6.** This figure illustrates a low confidence positive example (MT-GEM = 1) where two related molecules are incorrectly predicted by STEP. The likely reason for this is that the model failed to identify peak shifts characteristic of a single modification, giving little to no signal to predict MT-GEM 1, despite a high spectral similarity.

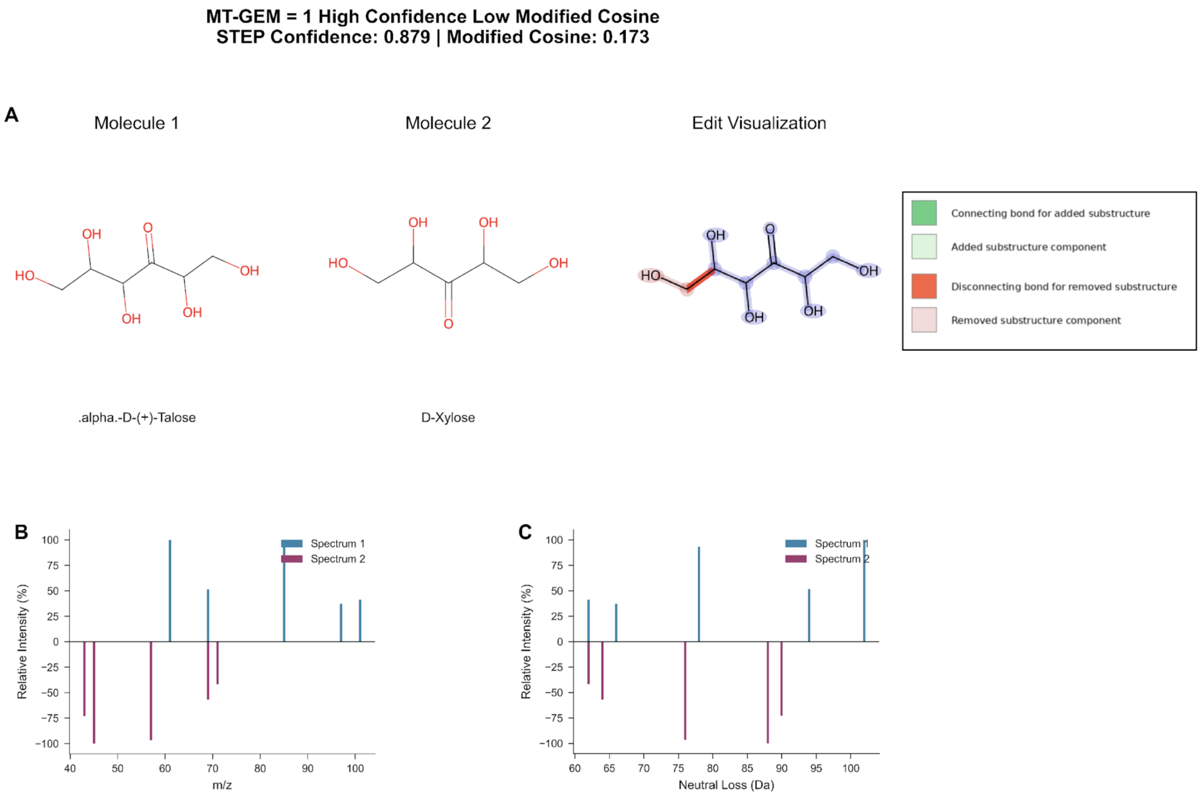


**Fig. S7.** This figure illustrates a high confidence positive example (MT-GEM = 1) where two related molecules are correctly predicted by STEP despite a low spectral similarity. The likely reason for this is that the model successfully identified MT-GEM 1 characteristic peak shifts that were not apparent in the modified cosine calculation.

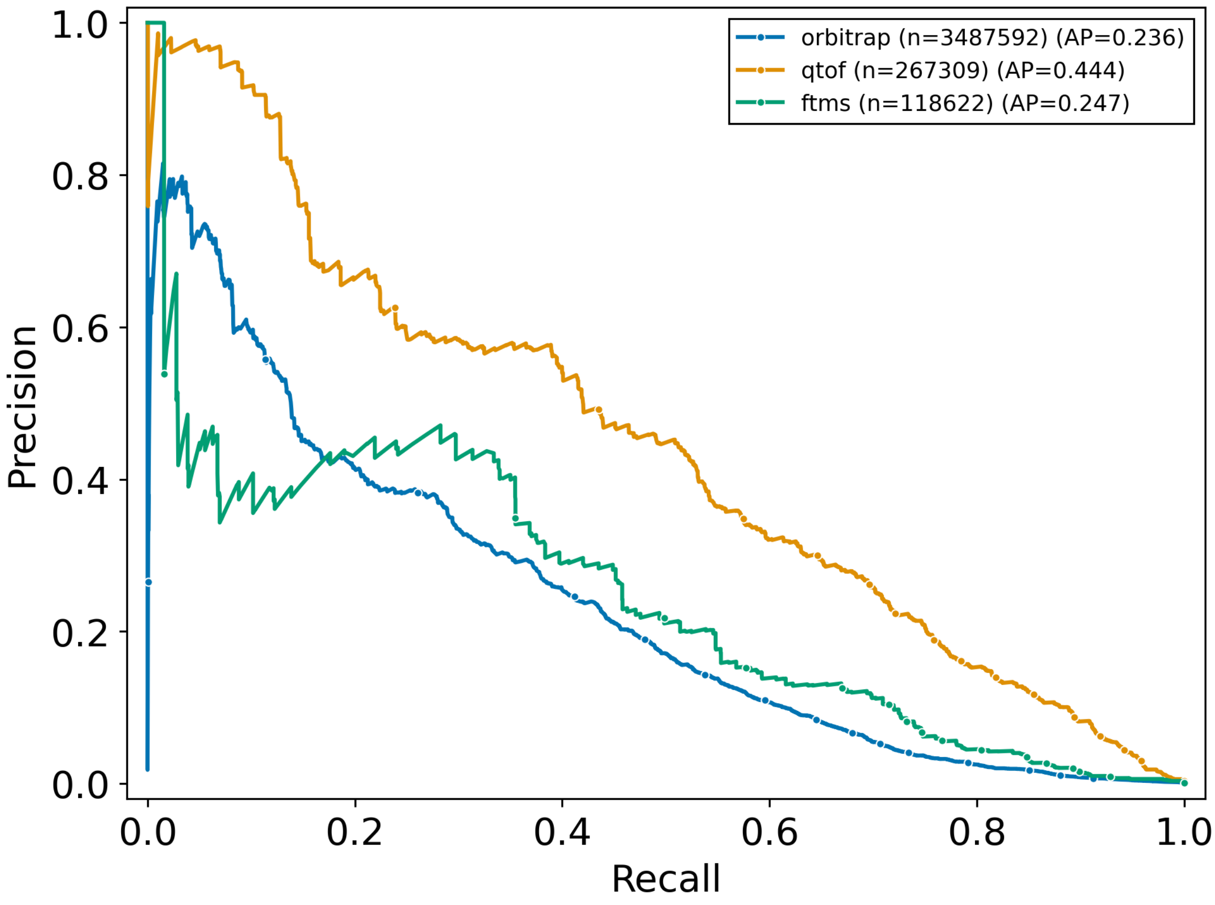


**Fig. S8.** Model performance across instrument types. Precision-recall curves for STEP evaluated on matched molecular structure sets with averaged precision/recall scores per unique structure pair across three mass spectrometry platforms. Sample sizes shown represent total spectral pairs per instrument type. QTOF instrument performance exceeds orbitrap and ftms performance. Performance variation likely reflects differences in spectral quality and fragmentation characteristics between instruments. Quadrupole data excluded due to limited sample size.

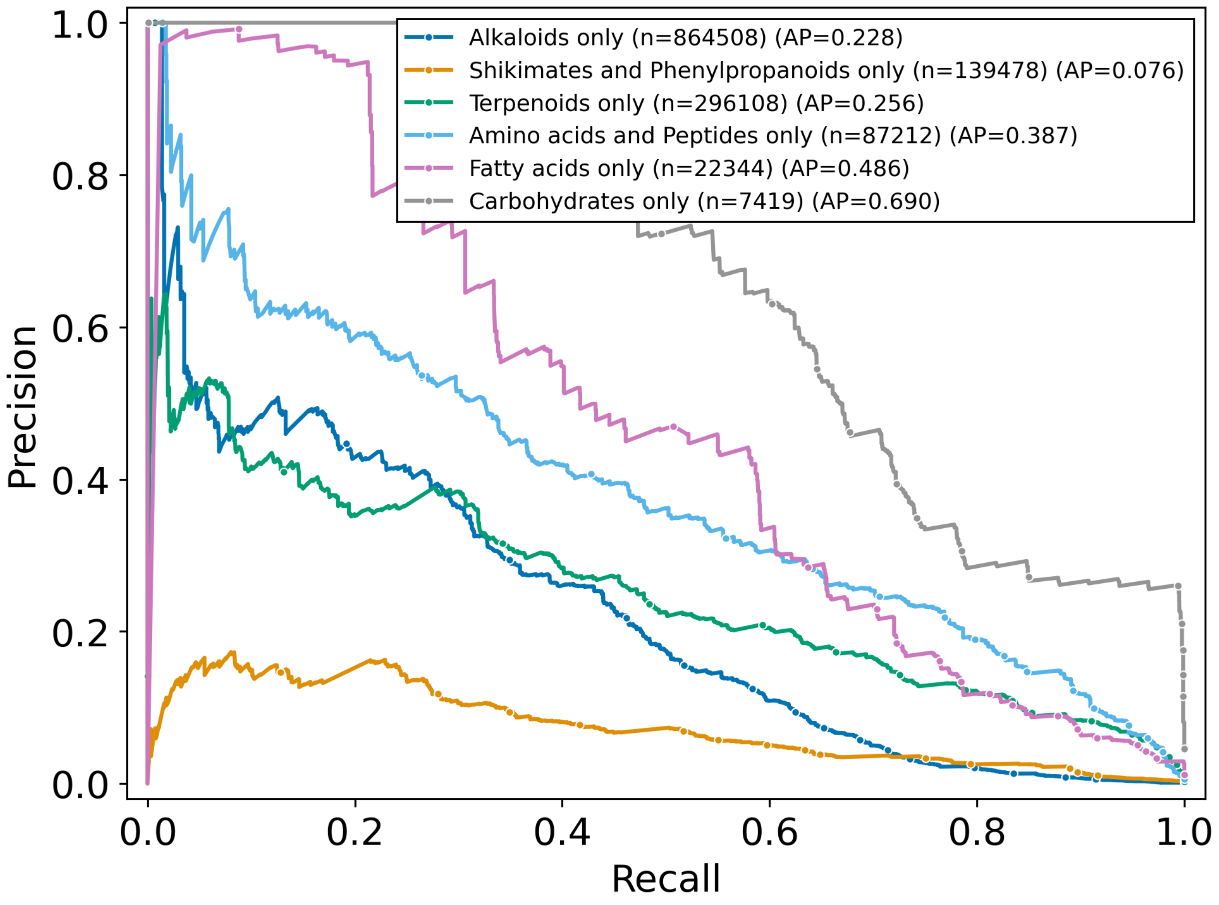


**Fig. S9.** STEP performance varies across different compound classes. Precision-recall curves with averaged precision/recall scores per unique structure pair for STEP evaluated on spectral pairs grouped by NPClassifier pathway annotations, highlighting large variation in performance for different compound classes. Pairs shown where both structures belong exclusively to the same compound class. Spectrum sample sizes shown in parentheses. Carbohydrate and fatty acid pairs exhibit notably superior performance, while shikimates and phenylpropanoid pairs show far lower average precision. Structures belonging to unknown compound classes were excluded.

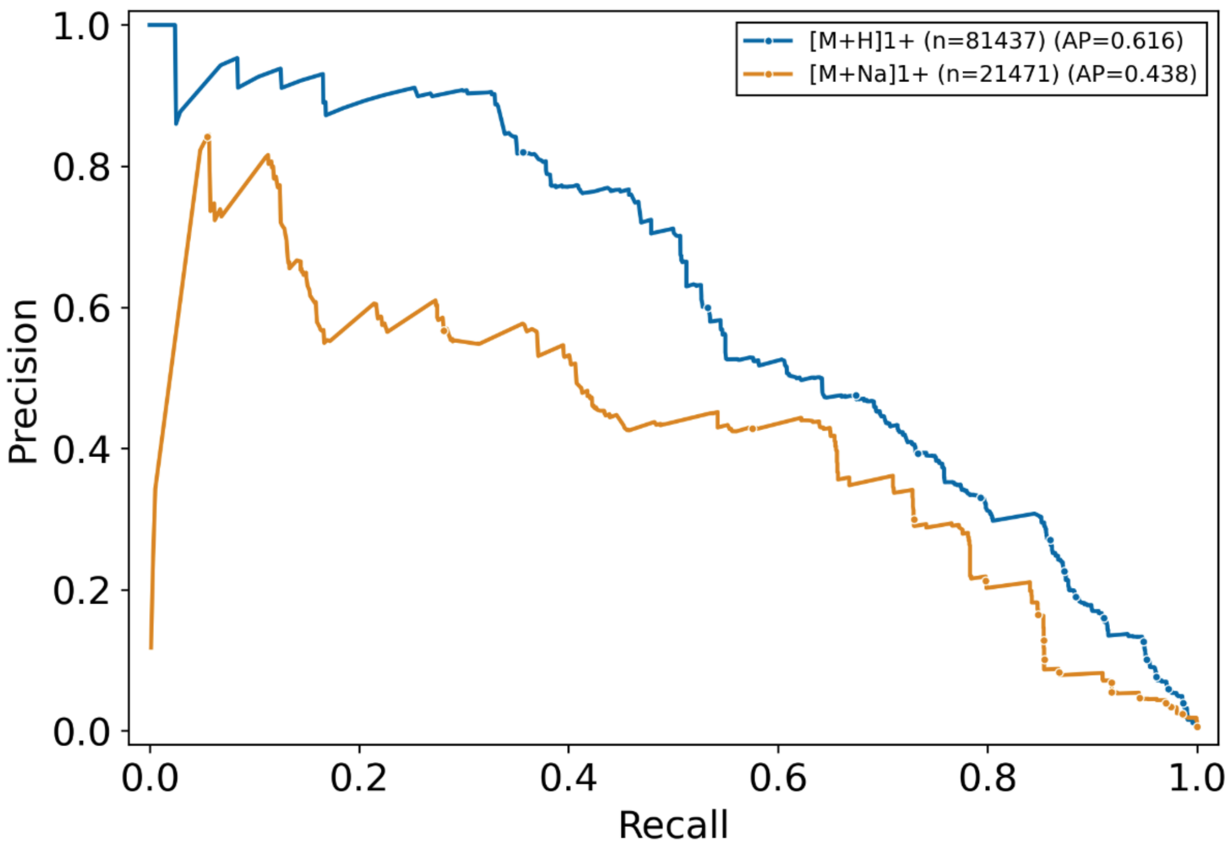


**Fig. S10.** Precision-recall curves for STEP evaluated on GNPSNIST test set pairs grouped by 2 most prevalent adduct types. We only show performance for structure pairs with +H and +Na adduct spectra, and average precision/recall performance per unique structure pair to control for imbalances in number of spectral pairs per structure and directly compare performance on different adducts. Hydrogen adducts are by far the most prevalent in library annotations, so the increased performance compared to sodium is expected. Likewise, The average precision for both curves appears to the average seen in Figure modelperformance, this is likely due to the fact that structures appearing with both +H and +Na adducts are likely overrepresented in library annotations due to their ease of annotation

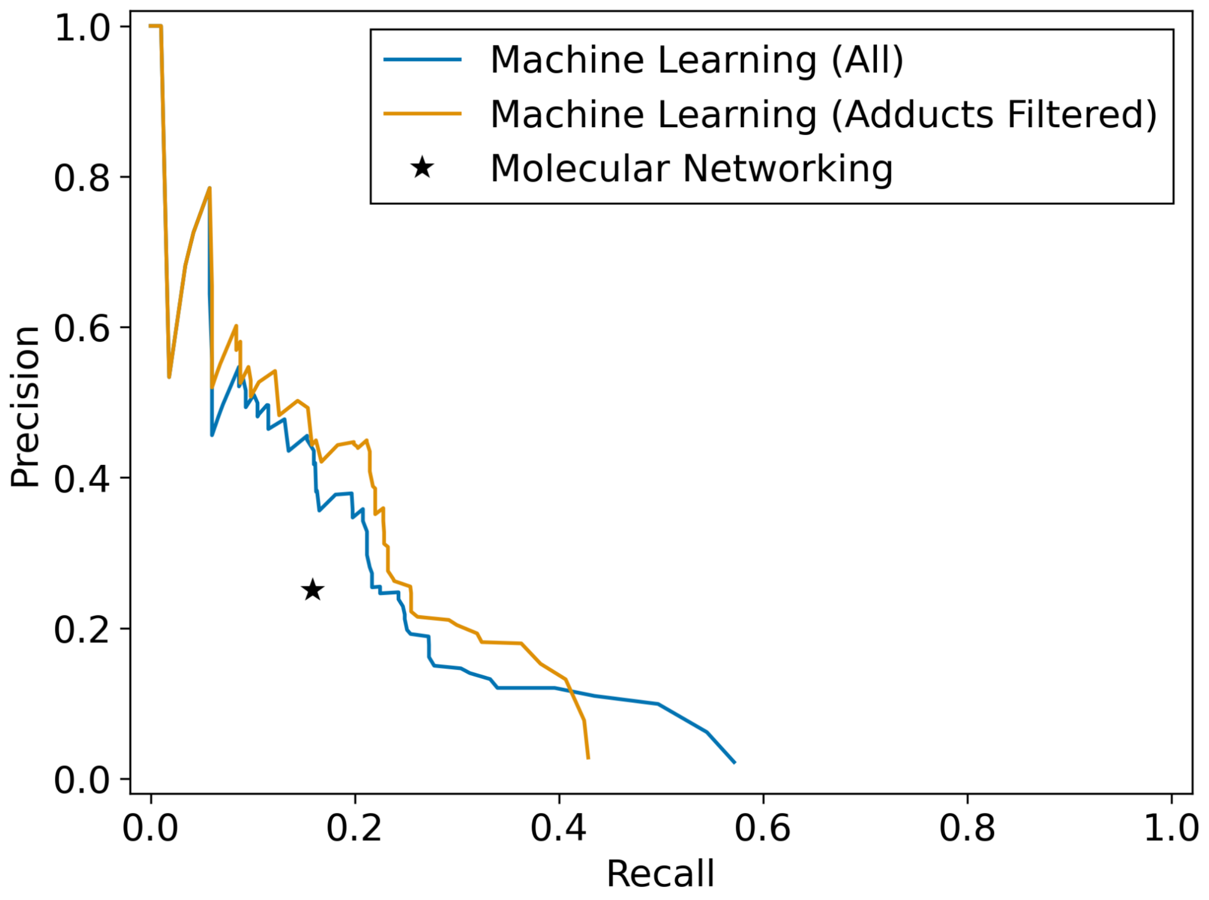


**Fig. S11.** Precision-recall curves for predicting MT-GEM distance 1 on the Com20 synthetic gut-community dataset (MSV000094899). We evaluate on library-matched spectra with known structures, considering only unique structure pairs. The blue line shows STEP performance without adduct filtering; the orange line shows performance when MS/MS pairs are required to share the same adduct. The star indicates feature-based molecular networking (FBMN) baseline performance (precision = 0.25, recall = 0.16, 40 unique SMILES pairs). At matched precision (0.25), STEP identifies approximately 1.5 times more single-transformation pairs than FBMN (recall 0.24 vs 0.16). At matched recall (0.16), STEP achieves 0.42 precision versus 0.25 for FBMN, a 1.67-fold improvement. Performance increases when MS/MS pairs are required to share the same adduct (see **Results MT-GEM Limitations**).

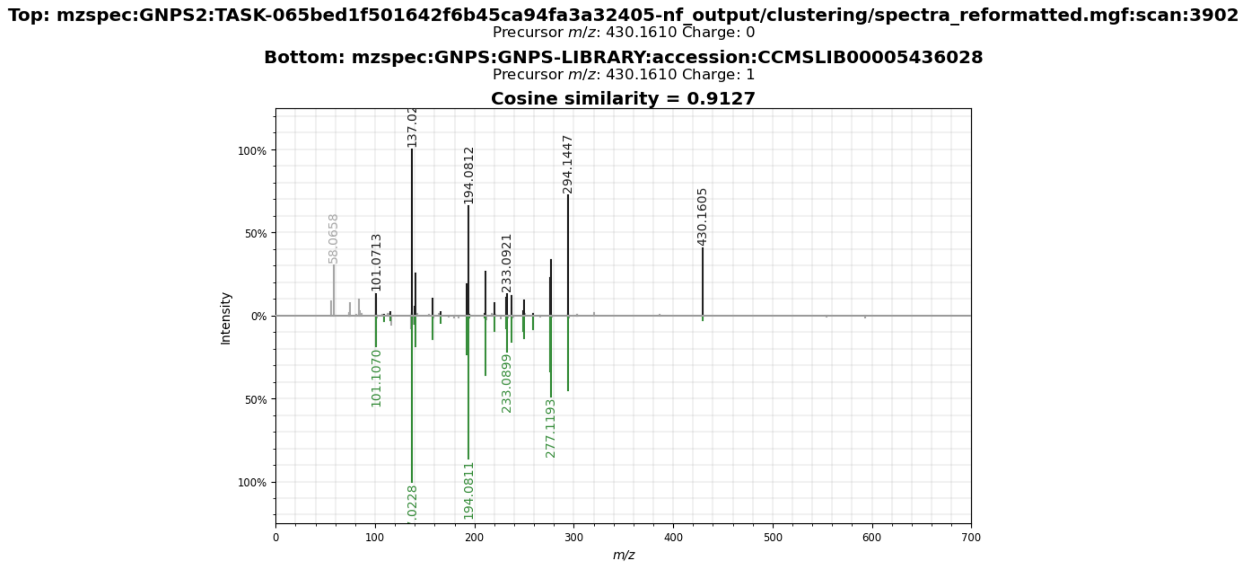


**Fig. S12.** Spectral library match supporting serratiochelin annotation. Mirror plot of the experimental MS/MS spectrum (top, precursor m/z 430.1610) acquired from the Serratia marcescens SPE fraction dataset against the GNPS library reference spectrum for serratiochelin (bottom, accession CCMSLIB00005436028, precursor m/z 430.1610). The match yields a cosine similarity of 0.91 (URL).

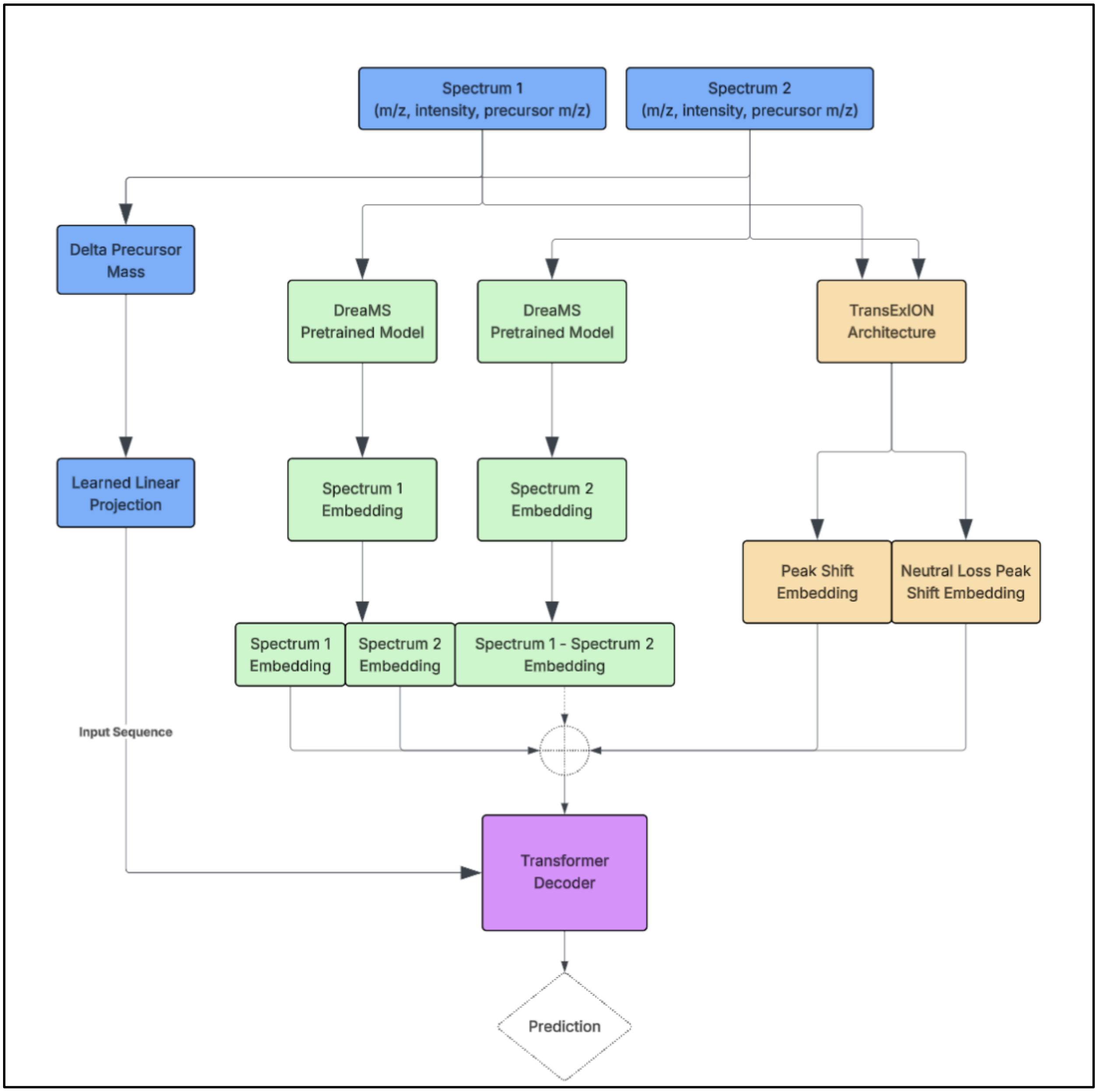


**Fig. S13.** STEP model architecture for predicting MT-GEM distance from MS/MS spectral pairs. Each input consists of two centroided MS/MS spectra with associated precursor m/z values. The precursor m/z values are first differenced to obtain the delta precursor mass, which is passed through a learned linear projection. In parallel, each spectrum is embedded using a DreaMS pretrained spectrum encoder to produce fixed-length spectrum embeddings; these are used individually (Spectrum 1 embedding, Spectrum 2 embedding) and as a difference vector (Spectrum 1 – Spectrum 2 embedding). Fragment-level relationships between the two spectra are captured by a TransExION-style peak-shift module, which encodes both direct peak shifts and neutral-loss peak shifts into two additional embeddings. All projected embeddings (delta precursor mass, three DreaMS-derived embeddings, and two peak-shift embeddings) are concatenated kids into an input sequence and fed to a transformer decoder, which outputs a final prediction, which is the probability that the spectrum pair corresponds to MT-GEM distance 1.

| Dataset | # Spectra | # Unique Structures |
|---|---|---|
| Train | 271,413 | 21,365 |
| Test | 51,183 | 3,740 |
| Validation | 6,306 | 500 |

Table 1 - Summary of the GNPSNIST dataset partitioning. Spectra were split into training, test, and validation sets. The validation set was randomly held out from the training set for hyperparameter tuning and early stopping.
